# The aPBP-type cell wall synthase PBP1b plays a specialized role in fortifying the *Escherichia coli* division site against osmotic rupture

**DOI:** 10.1101/2025.04.02.646830

**Authors:** Paula P. Navarro, Andrea Vettiger, Roman Hajdu, Virly Y. Ananda, Alejandro López-Tavares, Ernst W. Schmid, Johannes C. Walter, Martin Loose, Luke H. Chao, Thomas G. Bernhardt

## Abstract

A multi-protein system called the divisome promotes bacterial division. This apparatus synthesizes the peptidoglycan (PG) cell wall layer that forms the daughter cell poles and protects them from osmotic lysis. In the model Gram-negative bacterium *Escherichia coli*, PG synthases called class A penicillin-binding proteins (aPBPs) have been proposed to play crucial roles in division. However, there is limited experimental support for aPBPs playing a specialized role in division that is distinct from their general function in the expansion and fortification of the PG matrix. Here, we present *in situ* cryogenic electron tomography data indicating that the aPBP-type enzyme PBP1b is required to produce a wedge-like density of PG at the division site. Furthermore, atomic force and live cell microscopy showed that loss of this structure weakens the division site and renders it susceptible to lysis. Surprisingly, we found that the lipoprotein activator LpoB needed to promote the general function of PBP1b was not required for normal division site architecture or its integrity. Additionally, we show that of the two PBP1b isoforms produced in cells, it is the one with an extended cytoplasmic N-terminus that functions in division, likely via recruitment by the FtsA component of the divisome. Altogether, our results demonstrate that PBP1b plays a specialized, LpoB-independent role in *E. coli* cell division involving the biogenesis of a PG structure that prevents osmotic rupture. The conservation of aPBPs with extended cytoplasmic N-termini suggests that other Gram-negative bacteria may use similar mechanisms to reinforce their division site.

## Introduction

Cell division in bacteria is mediated by a multiprotein machine referred to as the divisome or septal ring^1^. This essential process is initiated by the coalescence of treadmilling polymers of the tubulin-like FtsZ protein into a dynamic ring pattern called the Z-ring at the prospective site of division^2–5^. The Z-ring is attached to the inner face of the cytoplasmic membrane via membrane-bound FtsZ interacting proteins like FtsA^6,7^. Following Z-ring formation, a number of essential and non-essential division proteins are recruited to the division site to form the mature divisome apparatus capable of promoting cytokinesis^1^.

A major function of the divisome is to synthesize the peptidoglycan (PG) cell wall material that will eventually fortify the poles of the daughter cells. Bacteria typically encode two main types of PG synthesis enzymes: (i) the bifunctional class A penicillin-binding proteins (aPBPs) that can both polymerize and crosslink glycans to form the PG matrix or (ii) the two-component synthases formed by complexes between a SEDS-family glycan polymerase and a monofunctional class B PBP (bPBP) with PG crosslinking activity^1^ (**Fig. 1A**). With only one known exception^8^, the essential bacterial cell division PG synthase is a SEDS-bPBP-type enzyme composed of the FtsW (SEDS) and FtsI (bPBP) proteins^9^. aPBPs have also been implicated in cell division, but their role in the process has not been clearly defined.

**Figure 1:**
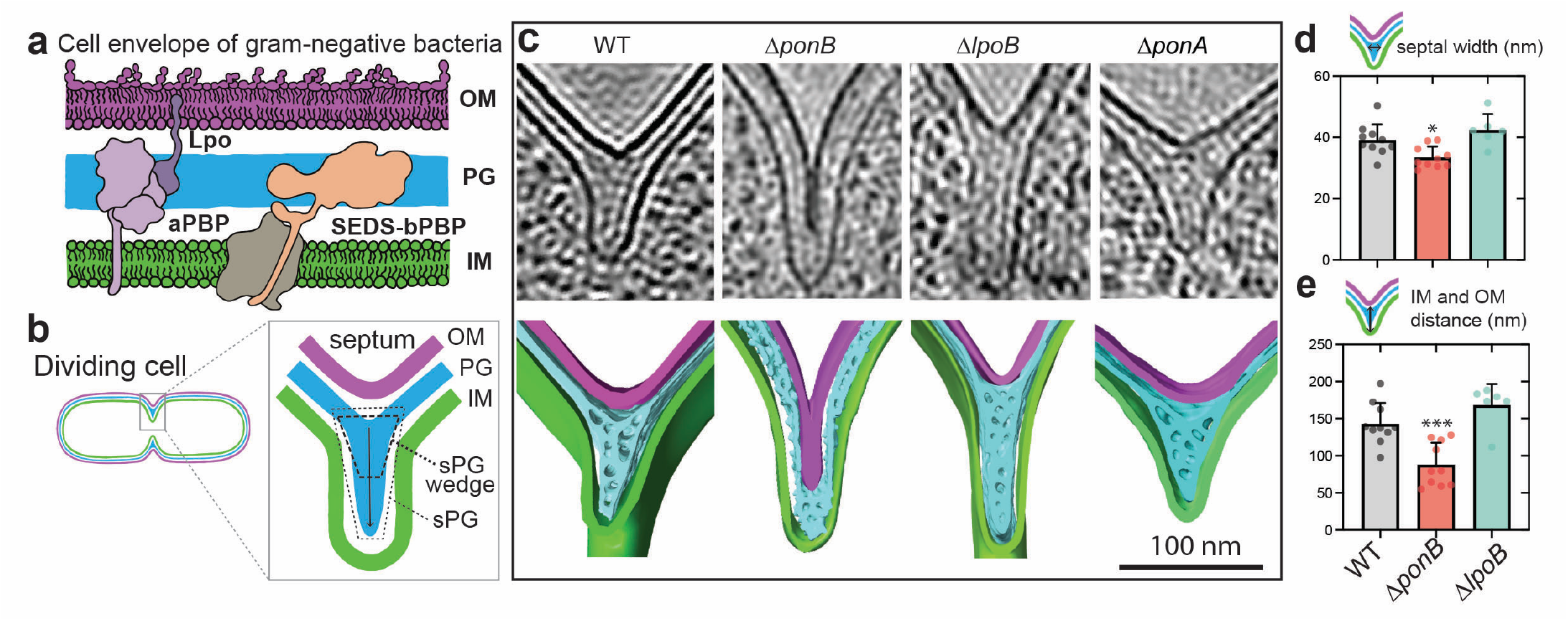
*In situ* septal PG architecture of wild-type and mutant *E. coli* cells. **(a)** Summary diagram of the different types of bacterial PG synthases. **(b)** Schematic showing the process of septum formation in Gram-negative bacteria. **(c)** Top row: summed, projected central slices of low-passed filtered cryo-electron tomograms visualizing sPG architecture of the indicated *E. coli* strains. Bottom row: 3D segmentations of the cryo-electron tomograms above rendering the IM as a green surface, the sPG as a cyan surface and the OM as a magenta surface. **(d)** Bar graph showing septal width measured orthogonal to the invagination direction in cryo-ET data (WT = 39.13 ± 5.13 nm; Δ*ponB* = 33.49 ± 3.5 nm; Δ*lpoB* = 42.43 ± 5.19 nm). Significance was tested using unpaired t-test with Welch correction for paired groups when data followed gaussian distribution and Mann-Whitney when data did not follow gaussian distribution and tested relative to wild-type. * = p < 0.05, ** = p < 0.01, *** = p < 0.001, **** = p < 0.0001. **(e)** Bar graph showing measured IM-OM distances at the septal region in the cryo-ET data. Thirty euclidean distances were measured per data point (see Methods), N values for each strain are: N = 10 (WT); 10 (Δ*ponB*); 6 (Δ*lpoB*). Scale bar = 100 nm.

*Escherichia coli* encodes two aPBPs that play important roles in PG biogenesis: PBP1a and PBP1b. These enzymes require a cognate outer membrane lipoprotein activator for their cellular activity^10,11^ (**Fig. 1A**). LpoA activates PBP1a whereas LpoB activates PBP1b (**Fig. 1A**). Neither enzyme system is essential, but their simultaneous inactivation is lethal^12,13^ and results in rapid cell lysis from lesions that develop at what appear to be random locations around the cell body as well as division site^10,11^. Thus, aPBPs are thought to play a general role in fortifying the PG matrix, especially in areas of low PG density or where damage has occurred^11,14,15^. The rate of PG synthesis in cells harboring PBP1b as the only aPBP is equivalent to that in WT cells and dramatically reduced when PBP1b is inactivated^14^. Additionally, mutants lacking PBP1b or LpoB, but not PBP1a or LpoA, are mechanically less stiff in the cylindrical region of the cell^16^ and are hypersensitive to beta-lactams and other perturbations to PG biogenesis^12,17–19^. Thus, of the two aPBP systems, PBP1b-LpoB appears to play the predominant role in the general growth and maintenance of the PG matrix.

In addition to a general role in the expansion and fortification of the PG matrix, PBP1b is also thought to play a significant role in cell division^20,21^. This functional assignment preceded the discovery of PG synthase activity for SEDS-bPBP complexes like FtsW-FtsI and thus was made essentially by default; the only choice before this time was one of the two aPBPs. Models that propose a role for PBP1b in division are principally based on protein crosslinking and co-purification experiments that identified interactions between PBP1b and several different components of the divisome^22–26^. What has been missing is a clear demonstration that these interactions and their effects on PBP1b activity observed *in vitro* are physiologically relevant and important for proper division in cells. Currently, the only genetic links between PBP1b and cell division are the observations that PBP1b inactivation results in a modest reduction in PG labeling at the division site^27^ and that deletion of the *ponB* gene encoding PBP1b is synthetically lethal with the inactivation of several non-essential division factors^28–30^. However, whether these phenotypes arise due to the general defect in PG biogenesis displayed by *ponB* mutants or a specific role for PBP1b in cell division is not known.

Here, we describe the discovery of a dedicated division function for PBP1b in *E. coli* enabled by our recent *in situ* ultrastructural analysis of *E. coli* division sites using cryo-focused ion beam (cryo-FIB) milling and cryogenic electron tomography (cryo-ET)^31^. Gram-negative bacteria like *E. coli* have a complex cell envelope made of two membranes with a relatively thin layer of PG cell wall sandwiched in the periplasmic space between them^32^ (**Fig. 1a**). When they divide, the inner membrane (IM) is invaginated and new PG material is synthesized within the invagination (**Fig. 1b**). Constriction of the IM at the division site is often observed to be deeper than that of the outer membrane (OM)^31,33,34^. Because this invaginating structure partially bisects the daughters, it is often referred to as the division septum and the associated cell wall material is called septal PG (sPG). This PG material is initially shared between daughter cells and must be progressively split during the division process by PG cleaving enzymes to allow the outer membrane to invaginate and eventually cover the new daughter cell pole^35^.

The combination of cryo-FIB milling and cryo-ET allowed us to resolve new features of septal architecture^31^. At the leading edge of the invaginating inner membrane, two plates of electron density were resolved that likely correspond to the PG material that will form the polar cell wall of the daughters. The architecture at the lagging edge of the septum closest to the outer membrane was markedly different. This region of the septum contained a thick, wedge-like density of PG^31^. The function of this so-called “sPG wedge” and the factors required for its synthesis have remained unclear. Given that genetic and imaging data point towards the FtsW-FtsI synthase functioning in sPG biogenesis at the leading edge of the invaginating septum^27,36^, we hypothesized that PBP1b might be functioning in the biogenesis of the sPG wedge. *In situ* cryo-ET imaging revealed that mutants lacking PBP1b indeed failed to form an observable sPG wedge. Additionally, atomic force microscopy (AFM) showed that PG purified from a mutant lacking PBP1b had a reduced stiffness at the division site relative to the sidewall whereas the opposite was true for PG isolated from wild-type cells. Notably, PBP1b inactivation was shown to result in a hypersensitivity to osmotic shock with the mutant cells lysing from lesions at midcell, indicative of a compromised septum. Surprisingly, the PBP1b activator LpoB^10,11^ was not required for sPG wedge formation or the osmotic stability of the septum. Finally, we show that of the two isoforms of PBP1b known to be produced in cells^37–39^, it is the longer protein with an extended cytoplasmic N-terminus that is recruited to the division site and that this recruitment is likely to be mediated by an interaction with the FtsA component of the Z-ring. Altogether, our results demonstrate that PBP1b plays a specialized, LpoB-independent role in *E. coli* cell division involving the biogenesis of a sPG wedge structure that fortifies the septum against osmotic rupture.

## Results

### PBP1b is required for sPG wedge formation

To investigate the potential role of PPB1b in sPG wedge formation we used *in situ* cryo-ET to image *E. coli* cells deleted for genes encoding PBP1b (Δ*ponB*) or its lipoprotein activator LpoB (Δ*lpoB*). Bacterial lamellae were generated by cryo-FIB milling, tilt-series were collected, and 3D reconstructed into cryo-electron tomograms. The tomograms were denoised and low-pass filtered to visualize sPG architecture *in situ* (**Table S1**). Densities corresponding to the IM (green), PG (blue), and OM (purple) were segmented, and template matching was used to localize oriented ribosome particles (gold) in the tomograms (see Methods; **Video S1-S4**). For comparison purposes, we used cryo-ET data of wild-type (WT) *E. coli* collected as part of our prior study^31^. Three dimensional (3D) renderings of the division site show architectural differences among strains (**Fig. S1**). A representative gallery of the cryo-ET data is available in **Fig. S2**. Cells lacking PBP1b had septa that were slightly thinner (16.8%) than those of WT cells when measuring the width of septa orthogonal to the direction of membrane invagination (**Fig. 1c-d**). Notably, however, the distance between the OM and IM within the constriction was significantly reduced in Δ*ponB* cells relative to WT with a corresponding loss of the wedge-like sPG density separating the two membranes in the cells lacking PBP1b (**Fig. 1c and 1e**). No notable differences in septal architecture were observed for cells lacking PBP1a versus WT (**Fig. 1c and S1**) as expected given that this aPBP has not previously been associated with major cell wall or division phenotypes^12,17–19^. Surprisingly, the architecture of septa in cells lacking the PBP1b activator LpoB was also largely comparable to that of WT, displaying similar width and OM-IM distance measurements (**Fig. 1c-e**).

To complement the cryo-ET imaging, we also examined the properties of sPG produced by WT and mutant cells by analyzing the height and stiffness of purified PG sacculi using atomic-force microscopy (AFM) (**Fig. 2a-d and Fig. S3**). Sacculi from WT cells had stiffer material at their division sites than material within the sidewall, but the opposite was true for sacculi isolated from cells lacking PBP1b, which had division sites with reduced stiffness relative to the sidewall (**Fig. 2d and S3a-b**). Another major difference between the PG preparations from WT and Δ*ponB* cells was in the frequency of lesions or “pores” in the PG network observed in the AFM scans for material thickness. Cells lacking PBP1b had an increased number of pores per total PG area and the pores that were observed in the Δ*ponB* sacculi tended to be larger in size that those detected in WT PG (**Fig. 2e-f and Fig. S3c**). This finding supports the idea that a major function of the PBP1b-LpoB system is to identify and “fill-in” areas of low density within the PG matrix^11,14,15^. Accordingly, PG sacculi isolated from Δ*lpoB* cells also displayed a similar increase in PG pore frequency to those isolated from cells lacking PBP1b (**Fig. 2d-f and Fig. S3**). However, despite the similar general PG biogenesis defect observed for Δ*ponB* and Δ*lpoB* sacculi, the sPG/side wall stiffness ratio was only reduced relative to WT for PG purified from cells inactivated for PBP1b not LpoB (**Fig. 2d**). Only a small reduction in the sPG/side wall stiffness ratio was observed for sacculi purified from cells lacking PBP1a (**Fig. 2d**). These sacculi also did not display a significant increase in the frequency of pores in the PG matrix (**Fig. 2d-f and Fig. S3**) as expected based on the lack of significant growth or antibiotic susceptibility phenotypes observed upon PBP1a inactivation. Based on the combination of the cryo-ET and AFM analyses, we conclude that PBP1b is required for the biogenesis of the sPG wedge structure at the division site. Furthermore, this division function is not dependent on the LpoB activator, indicating that it is a specialized activity of the PG synthase distinct from its general role in PG biogenesis.

**Figure 2:**
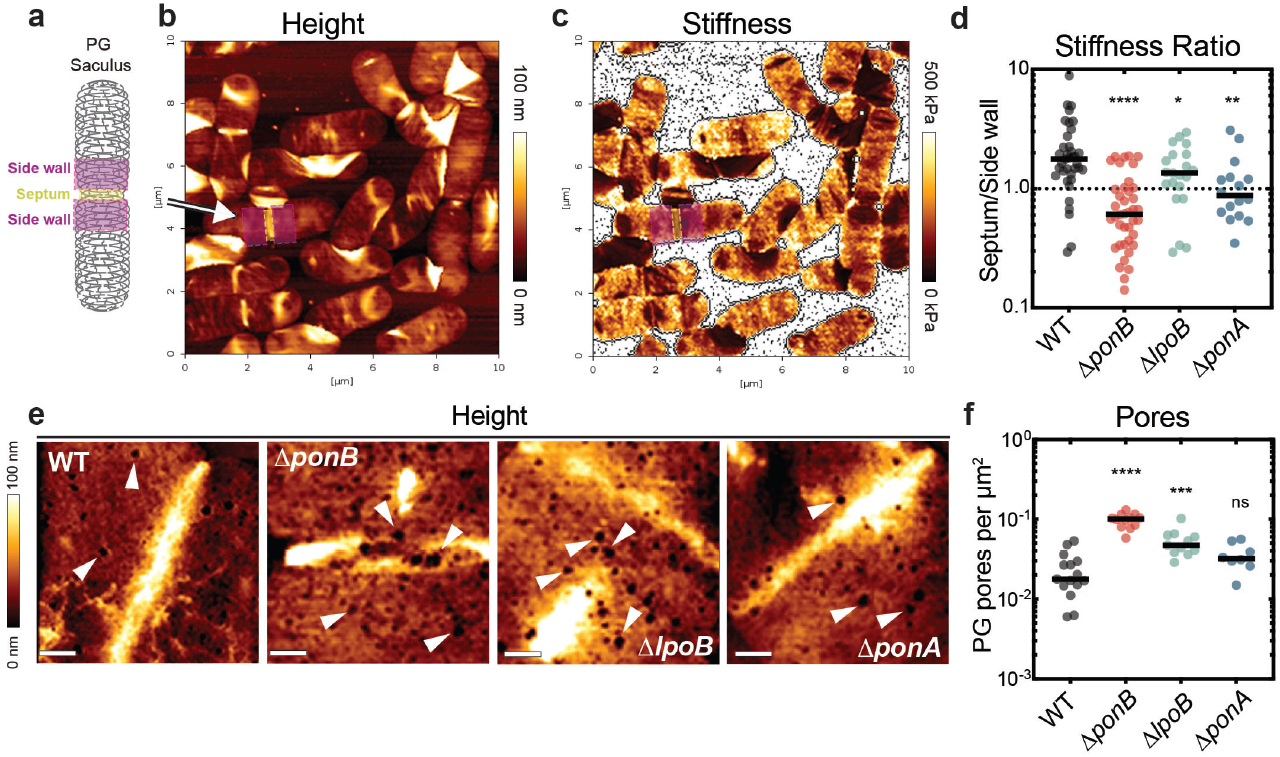
AFM analysis of PG sacculi from wild-type and mutant *E. coli* cells. PG sacculi from the indicated strains were isolated and imaged by AFM. **(a)** Schematic overview of septal (yellow box) and sidewall (magenta box) regions analyzed for stiffness. Representative images from (b)height and **(c)** stiffness (Young’s moduli) scans of WT PG sacculi at low resolution (10 × 10 µm, 39.1 nm pixel size). Regions of analysis for septal/sidewall comparisons are highlighted with an arrow and colored as in **(a). (d)** Ratio of septal to side wall PG stiffness was measured for the indicated strains and significant differences relative to WT were determined using one-way ANOVA with Dunnett’s posttest. * = p < 0.05, ** = p < 0.01, **** = p < 0.0001, N sacculi for each strain from 3 biological replicates were measured: N = 37 (WT); 40 (Δ*ponB*); 21 (Δ*lpoB*); 15 (Δ*ponA*). **(e)** Representative high-magnification (1 × 1 µm, 7.81nm pixel size) height images of septa isolated from indicated strains. Arrow heads point to pores identified in the cell wall. Scale bar = 200 nm. **(f)** Quantification of pores per µm^2^. Significant differences relative to WT were determined using one-way ANOVA with Dunnett’s posttest. *** = p < 0.001, **** = p < 0.0001, ns = non-significant, N sacculi for each strain from 3 biological replicates were measured: N = 15 (WT); 12 (Δ*ponB*); 11 (Δ*lpoB*); 8(Δ*ponA*).

### The sPG wedge is required to fortify the division site against osmotic rupture

We previously speculated that the sPG wedge was likely to be important for the osmotic stability of the septum^31^. Consistent with this hypothesis, mutants inactivated for PBP1b were found to have a severe growth defect at high temperature (42°C) in medium with a low osmolyte concentration (half-strength LB medium with no added NaCl, 0.5xLB0N) (**Fig. 3a**). A mutant defective for LpoB grew normally under this condition, indicating that the phenotype is not due to general cell wall defects (**Fig. 3a**). Growth was also unaffected by PBP1a inactivation (**Fig. 3a**). Thus, a defect in sPG wedge formation compromises growth in an osmotically challenging condition.

**Figure 3:**
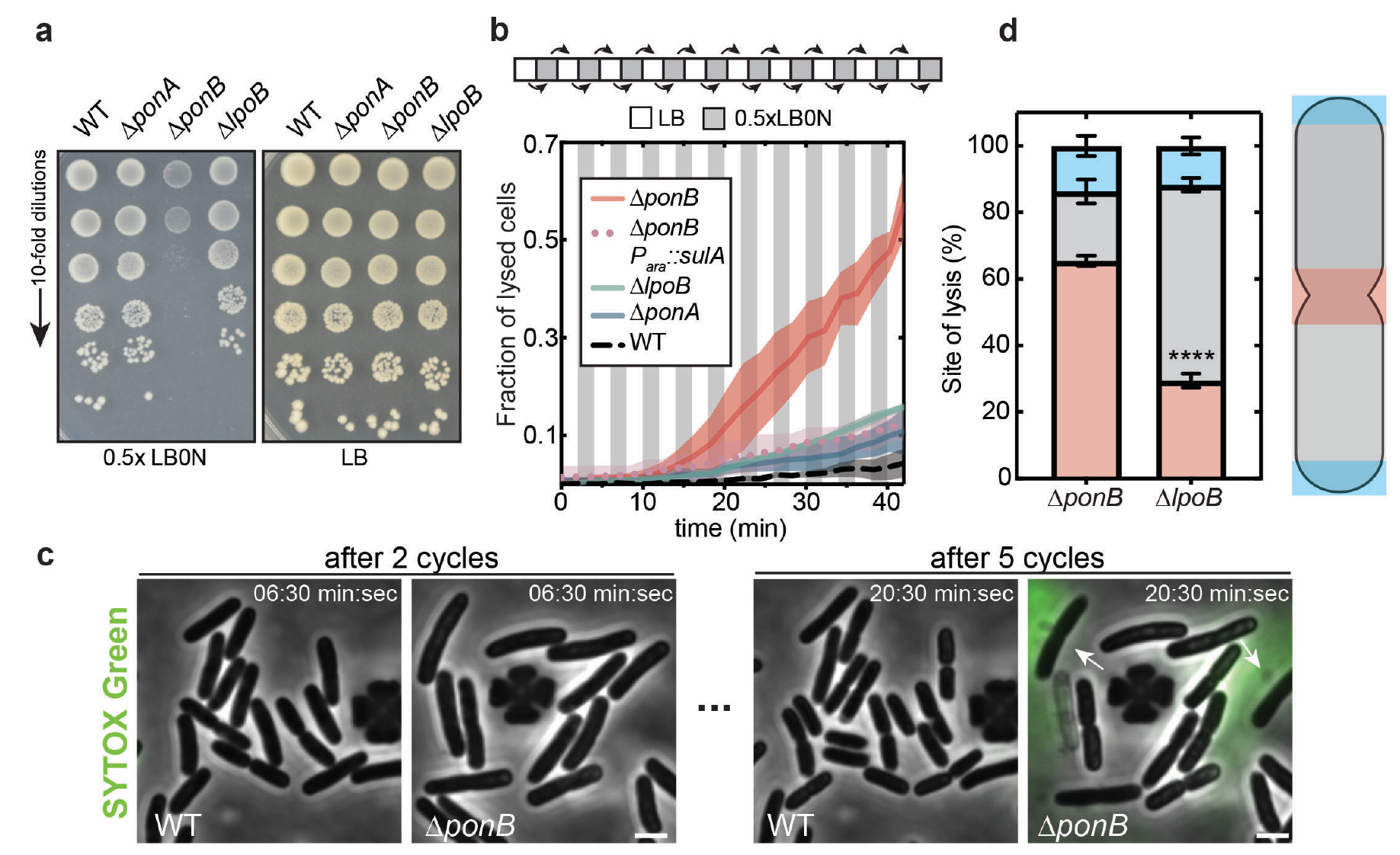
PBP1b stabilizes the *E. coli* division site against osmotic rupture. **(a)** Representative image of a bacterial viability assay. Cells were grown on LB or 0.5xLB0N agar at 42°C as indicated. **(b)** Osmotic oscillations between LB (white) and 0.5xLB0N (grey) were performed using the CellAsic microfluidic platform. The fate of cells from the indicated strains was monitored at 0.1 Hz for 42min by phase contrast and SytoxGreen (1 µM) imaging. Filamentation was induced by expression the division inhibitor SulA for 20 min prior to exposure to the first osmotic shock. SulA production was induced using 0.2% L-arabinose. Graph shows the mean fraction of lysed cells (line) ± one SD (shade) plotted from N biological replicates: N = 6 (WT); 5 (Δ*ponB*); 4 (Δ*lpoB*); 3 (Δ*ponB* +SulA); 3 (Δ*ponA*). **(c)** Representative images of indicated strains after two and five cycles of osmotic shocks. Arrow point towards sites of septal lysis events. Scale bar = 2 µm. **(d)** Manual quantification of subcellular lysis sites in response to osmotic shocks from 100 cells from 3 biological replicates. Data is represented as mean ± one SD and significance was tested using unpaired t-test, **** = p < 0.0001.

To investigate the nature of the osmotic sensitivity displayed by Δ*ponB* cells and whether it was the result of a failure in septal integrity, we performed a time-lapse imaging experiment that followed cell growth and morphology in a microfluidic chamber during cycles of osmotic shifts between normal LB and 0.5xLB0N (**Fig. 3b-d and Video S5**). Imaging was performed at 37°C where the Δ*ponB* growth defect is less severe than at 42°C. Sytox Green was also included in the medium to stain extracellular DNA and aid the visualization of cell lysis. Over a series of twenty osmotic shifts, WT cells grew normally with minimal lysis events observed (4.44 ± 3.07% lysed cells) (**Fig. 3b-c and Video S5**). A slight elevation in the frequency of cell lysis was observed for cells lacking LpoB (15.92 ± 8.96% lysed cells) or PBP1a (11.01 ± 5.12 % lysed cells), but lysis was dramatically increased in cells inactivated for PBP1b (58.81 ± 7.51 % lysed cells) (**Fig. 3b-c, S4 and Video S5**). Importantly, blocking cell division by expression of the FtsZ antagonist SulA greatly reduced the level of lysis observed for Δ*ponB* cells (12.51 ± 3.54 % lysed cells) (**Fig. 3b-c, S4, and Video S5**). Moreover, when the site of cell lysis was analyzed based on the location of membrane bleb formation and DNA release, cells lacking PBP1b lysed from septal failures at a much higher rate than cells lacking LpoB (**Fig. 3d**). These results indicate that PBP1b is critical for the stabilization of the division septum likely via its role in sPG wedge formation.

### The alpha isoform of PBP1b is enriched at the division site

Our results thus far suggest that PBP1b plays a specialized role in cell division distinct from its general function in PG biogenesis. Such a role would suggest that the enzyme is recruited to the division site to participate in the process. However, PBP1b has not been found to be strongly enriched at the division site. Immunofluorescence microscopy has previously observed a weak enrichment of PBP1b at midcell relative to the periphery in normally growing cells^22,40^. The enrichment is more pronounced in cells treated with the beta-lactam aztreonam^25^, but whether this localization reflects a division activity or a more general repair activity in response to PG damage caused by the division-specific PG synthesis inhibitor is not known. Furthermore, unlike the immunofluorescence results, green fluorescent protein (GFP) fusions to PBP1b displayed a peripheral localization pattern without a discernable enrichment at the division site^10^. Thus, whether PBP1b is specifically recruited to the septum to function in division has remained unclear.

We wondered whether the difficulties defining the localization of PBP1b and its importance for cell division might be related to the observation that *E. coli* cells produce two isoforms of the enzyme^37–39^. The alpha form of PBP1b (^α^PBP1b) has a longer cytoplasmic N-terminal domain than the gamma form (^γ^PBP1b) for which translation is initiated 46 codons downstream of the ^α^PBP1b start (**Fig. 4a**). To determine if the two isoforms have differential subcellular localization patterns, cells producing monomeric superfolder GFP (msfGFP) fusions to either ^α^PBP1b (Met 46 changed to Leu) or ^γ^PBP1b were imaged. When either fusion was produced at low levels of induction (50 µM IPTG), newly born cells without an observable midcell constriction displayed a weak peripheral fluorescence signal indicative of a dispersed localization throughout the membrane (**Fig. 4b-c**). In longer cells with a constriction, only the msfGFP-^α^PBP1b fusion appeared moderately enriched at the division site relative to the peripheral membrane (**Fig. 4b-c**). On the other hand, the shorter msfGFP-^γ^PBP1b fusion maintained its largely peripheral pattern with weak midcell enrichment that is likely a consequence of membrane invagination causing an increase in the membrane signal (**Fig. 4b-c**). These localization patterns were observable in single cells but were most clear upon analysis of cell populations using demographs, which show fluorescence intensity profiles across the bodies of hundreds of cells. When these profiles are stacked according to cell length to create the demograph, patterns of protein localization as a function of the cell cycle become evident. For cells producing msfGFP-^α^PBP1b, an increase in fluorescence intensity was observable at midcell in the population of longer (older) cells (**Fig. 4b**). This localization appeared to be roughly coincident with the onset of cell constriction observed as a decrease in midcell width in this population when the corresponding phase contrast images were analyzed (**Fig. 4b**). When analyzed similarly, cells producing msfGFP-^γ^PBP1b did not display nearly as strong of a midcell enrichment as those producing msfGFP-^α^PBP1b (**Fig. 4b-c**), suggesting that the extended N-terminus of the alpha isoform is required for recruitment of PBP1b to the division site.

**Figure 4:**
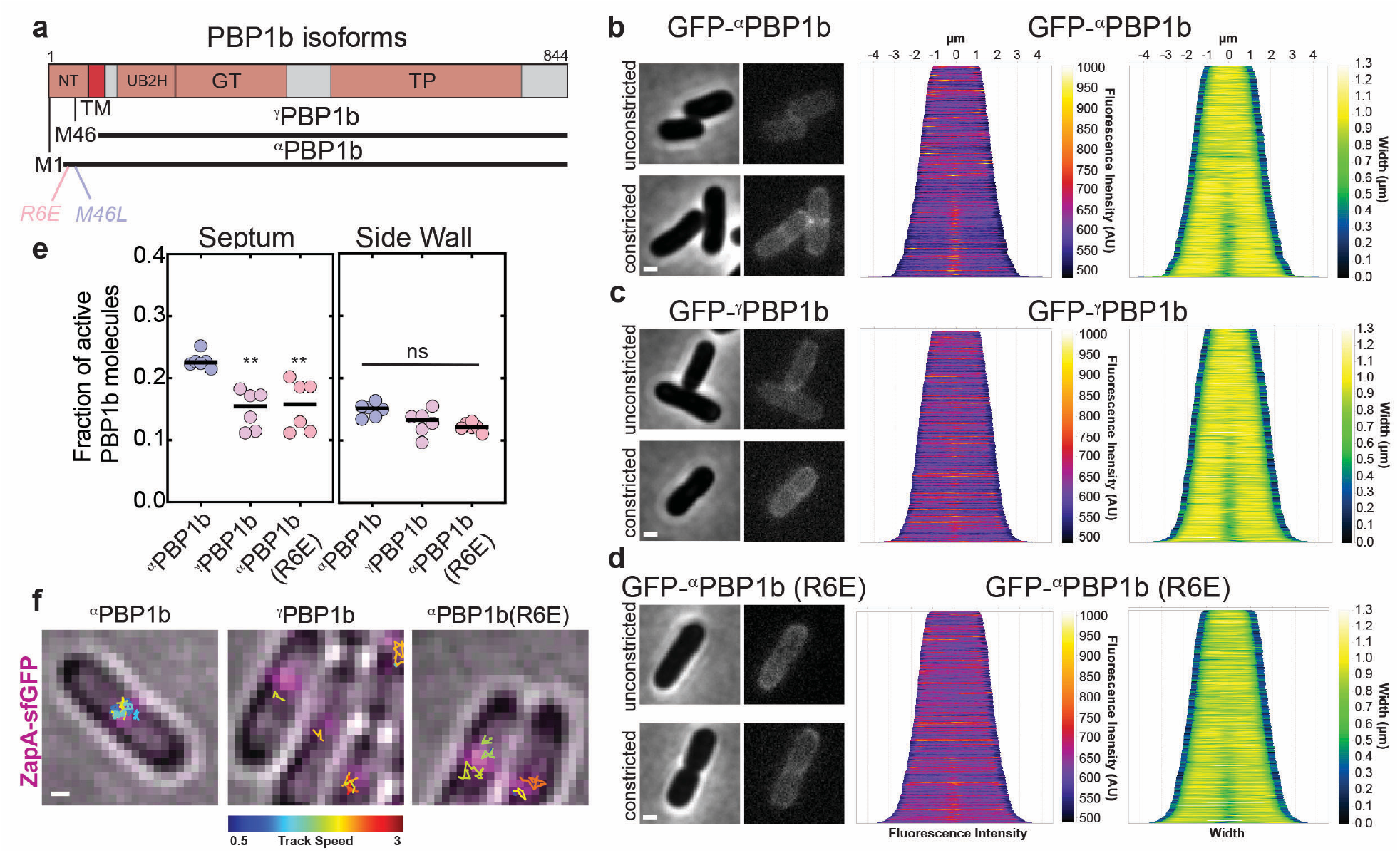
The longer alpha isoform of PBP1b is recruited to the division site. **(a)** Primary structure, domain architecture, and isoforms of PBP1b. NT = cytoplasmic N-terminus, TM = transmembrane helix, UB2H = LpoB binding domain, GT = PG polymerase domain, TP = transpeptidase domain. **(b-d)** (Middle) Demographs (N = 395 (GFP-^α^PBP1b); 401 (GFP-^γ^PBP1b); 629 (GFP-^α^PBP1b(R6E)) showing localization of the indicated GFP-PBP1b isoform across the cell population arranged by increasing cell length (cell age). (Right) Demographs of cell width measurements across their length. (Left) Representative images of cells with or without an observable septal invagination are shown on the left. Demographs and snapshots are representative from 3 biological replicates. **(e)** Mean of active fraction (stationary particles) of indicated PBP1b molecules as determined from SPT of Halo-PBP1b fusions acquired at 20 Hz. Significance was determined using one-way ANOVA with Tucky’s posttest. ** = p < 0.01, ns = non-significant. N = 6 SPT experiments from two biological replicates. **(f)** Representative images of SPT tracking data for Halo-PBPB1b isoforms. Septal PBP1b SPT trajectories are overlayed onto composite of 2×2 binned bright field and GFP reference image. ZapA-GFP (false-colored in magenta) signal served as a marker for division site localization. Scale bars = 0.5 µm.

Prior analyses of aPBP dynamics detected two distinct populations of molecules, diffusive and stationary, with the stationary (bound) molecules being associated with PG synthesis function^14,15^. We therefore compared the single molecule dynamics of Halo fusions to the two PBP1b isoforms to test for potential differences in activity at the division site using total internal reflection microscopy (TIRFM) and single particle tracking (SPT). Obtained tracks were fitted to a two-state kinetic model to determine the fraction of bound molecules^41–43^. To analyze the displacement of single PBP1b molecules on or in close vicinity to the Z-ring, ZapA-sfGFP was used as a fiducial marker. A similar fraction of active (i.e. stationary/bound) molecules were detected along the sidewall for both the Halo-^α^PBP1b and Halo-^γ^PBP1b fusions used for the tracking analysis (**Fig. 4e, S5a, and Video S6**). However, when molecules in the region of the division site were analyzed, a higher proportion of the Halo-^α^PBP1b molecules were active than those of Halo-^γ^PBP1b (**Fig. 4e-f, S5b, and Video S6**). Thus, these imaging experiments indicate that the longer N-terminal domain of the ^α^PBP1b isoform is associated with better midcell recruitment and higher activity at the division site.

### Evidence that αPBP1b is recruited to the division site via an interaction with FtsA

To identify potential partners of the ^α^PBP1b isoform that promote its recruitment to the division site via interaction with its N-terminal cytoplasmic domain, we performed an *in silico* protein-protein interaction screen^44^. Specifically, we used AlphaFold-Multimer to “fold” an N-terminal peptide of ^α^PBP1b or an N-terminal peptide of ^γ^PBP1b with all ∼4200 *E*.*coli* proteins, and we scored these prediction using several metrics (see methods). In total, 579 significant hits were obtained (^α^PBP1b: 442; ^γ^PBP1b: 137) (Extended Data Table 1). Among the top hits, many likely correspond to false positives given that they reside in the wrong cellular compartment (e.g. periplasm and OM). Nevertheless, we were excited to see that the divisome protein FtsA was among the potential interaction partners of the N-terminal peptide of ^α^PBP1b but not of ^γ^PBP1b. The structural prediction suggests that the N-terminal peptide of ^α^PBP1b, referred to as ^N-pep^PBP1b, might bind FtsA with an interaction involving a salt-bridge between R6 of ^α^PBP1b and E303 of FtsA (**Fig. 5a**). Notably, this region of FtsA is where the C-terminal peptide of FtsZ is bound to promote the association of FtsZ polymers with the inner face of the IM, suggesting that the N-terminal domain of PBP1b may compete with FtsZ for binding to FtsA (**Fig. S6**).

**Figure 5:**
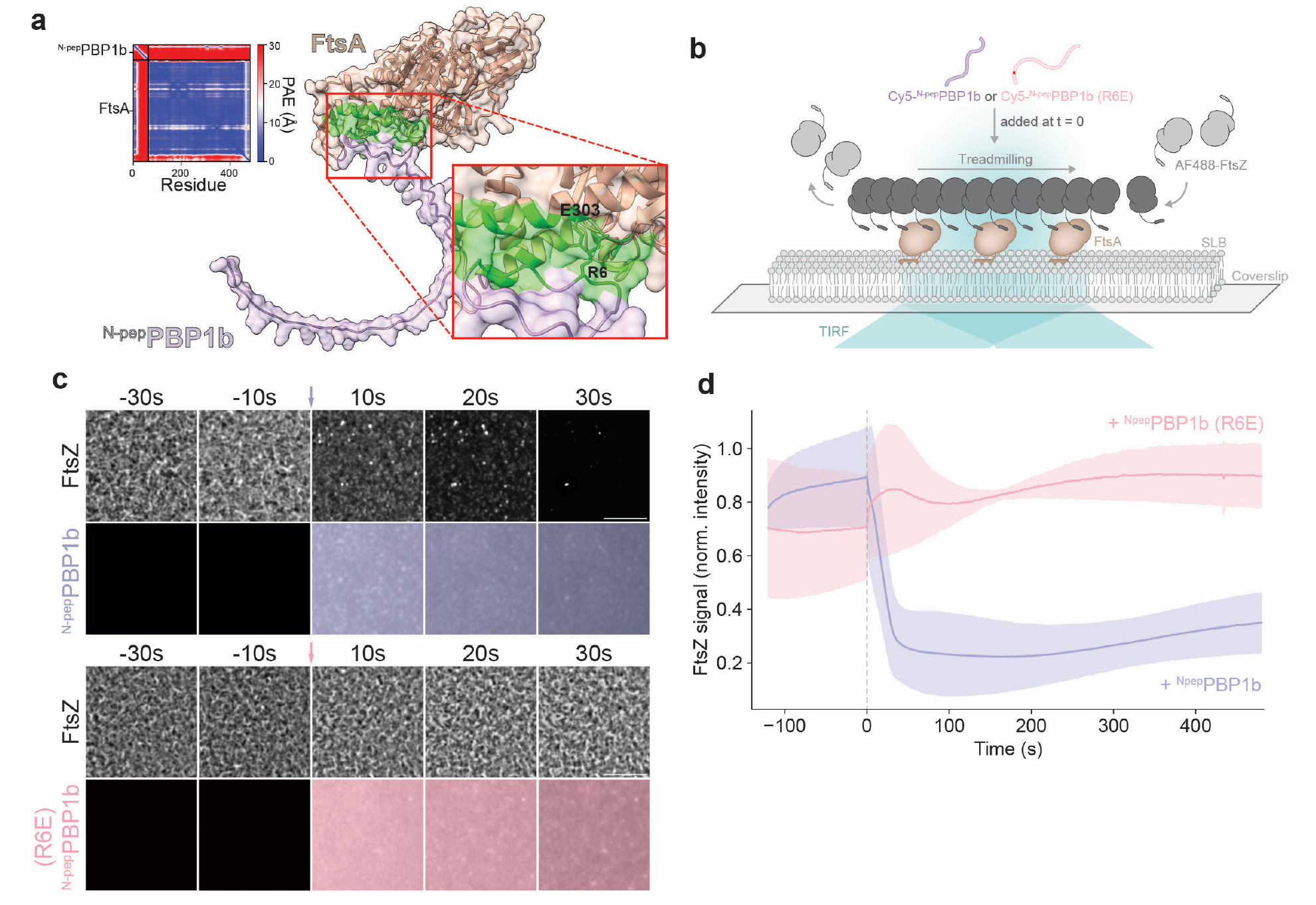
Evidence that the N-terminal domain of αPBP1b interacts with FtsA. **(a)** Predicted structure FtsA (salmon) with ^N-pep^PBP1b (purple) with the salt-bridge (^α^PBP1b(R6)-FtsA(E303)) at the interaction interface highlighted in green. Red box indicates the magnified region shown on the right. Predicted alignment error in Å (top left) of all residues against all residues for the top-ranked model (right). Low error (blue) corresponds to well-defined relative domain positions. Right: Magnification showing the interaction between FtsA-E303 and ^α^PBP1b-R6 (green). **(b)** Sideview schematic portraying experimental setup and protein components for experiments in panels c and d. **(c)** Representative micrographs of AF488-FtsZ (grey), Cy5-N-^α^PBP1b (purple) and Cy5-N-^α^PBP1b (R6E) (pink). Arrows (t = 0 s) correspond to the time of addition of corresponding peptide. Scale bar = 5 µm. **(d)** Mean intensity ± one SD (shade) over time projections of normalized FtsZ fluorescence signal at the membrane before and after addition (t = 0 s) of Cy5-N-pepPBP1b (purple) and Cy5-N-pepPBP1b (R6E) (pink) plotted from 3 technical replicates.

Numerous attempts to detect a direct interaction between FtsA and ^N-pep^PBP1b using pull-downs with purified components or bacterial two-hybrid assays were unsuccessful. However, these assays did not capture the cellular context of the interaction, which likely involves polymeric forms of FtsA associated with the membrane where ^α^PBP1b is integrated. Reasoning that such a context may be needed to promote the putative interaction via avidity effects and an increase in effective protein concentration at the membrane, we turned to a previously developed assay for monitoring the recruitment of FtsZ protofilaments to supported lipid bilayers by FtsA polymers^5^ (**Fig. 5b**). In this assay, lipid bilayers are formed on coverslips and purified FtsA and FtsZ proteins are added to the solution above the membrane. The FtsZ molecules are fluorescently labeled and their localization and dynamics on the membrane are monitored by TIRFM. Importantly, FtsZ localization to the membrane in this assay is dependent on its interaction with FtsA^5^. Given that ^N-pep^PBP1b is predicted to compete with FtsZ for binding to FtsA, we used FtsZ membrane association as a proxy for the detection of a potential ^N-pep^PBP1b-FtsA interaction. Treadmilling FtsZ polymers were formed in the presence of FtsA on membrane bilayers doped with Ni-NTA-containing lipids. Upon the addition of His-tagged ^N-pep^PBP1b, the FtsZ polymers briefly colocalized with ^N-pep^PBP1b (**Fig. S7**) and then were rapidly dissociated from the membrane (**Fig. 5c-d, Video S7**). However, FtsZ polymers at the membrane were largely unaffected by the addition of His-tagged ^N-pep^PBP1b(R6E) (**Fig. 5c-d, Video S8**), which harbors a substitution predicted to disrupt the ^N-pep^PBP1b-FtsA interaction. Thus, the FtsZ membrane recruitment assay supports the AlphaFold2 prediction that ^N-pep^PBP1b binds FtsA in its FtsZ-binding region and that the salt bridge between R6 of ^α^PBP1b and E303 of FtsA plays an important role in this interaction.

### Midcell recruitment of αPBP1b is critical for its role in fortifying the septum

To investigate the role of the predicted ^α^PBP1b-FtsA interaction on the recruitment of PBP1b to the division site, we compared the subcellular localization of GFP-^α^PBP1b(WT) and GFP-^α^PBP1b(R6E). Similar to GFP-^γ^PBP1b, GFP-^α^PBP1b(R6E) also was not strongly recruited to midcell (**Fig. 4b-d**). Additionally, single particle tracking of the corresponding Halo fusions showed that a significantly (p = 0.0024) higher proportion of ^α^PBP1b(WT) was active at the division site relative to ^α^PBP1b(R6E) (**Fig. 4e-f, Fig S5a-b, Video S6**). We next tested the functionality of ^α^PBP1b(R6E) relative to ^α^PBP1b(WT) based on their ability to complement the midcell lysis defect of Δ*ponB* cells observed following osmotic oscillations between LB and 0.5xLB0N. As expected, Δ*ponB* cells producing ^α^PBP1b(WT) were largely resistant to lysis induced by the osmotic changes (**Fig. 6a-b, Video S5**). Intriguingly, cells expressing the ^α^PBP1b(R6E) showed the same increase in septal lesions as cells harboring the full deletion of PBP1b (**Fig. 6c**). However, despite its accumulation to similar levels in cells (**Fig. S8**), ^α^PBP1b(R6E) production failed to suppress the midcell lysis phenotype caused by the deletion of *ponB* (**Fig. 6c**). Based on these results, we conclude that the recruitment of ^α^PBP1b to the division site is required for its function and that this recruitment is likely to be mediated by the predicted ^N-pep^PBP1b-FtsA interaction.

**Figure 6.**
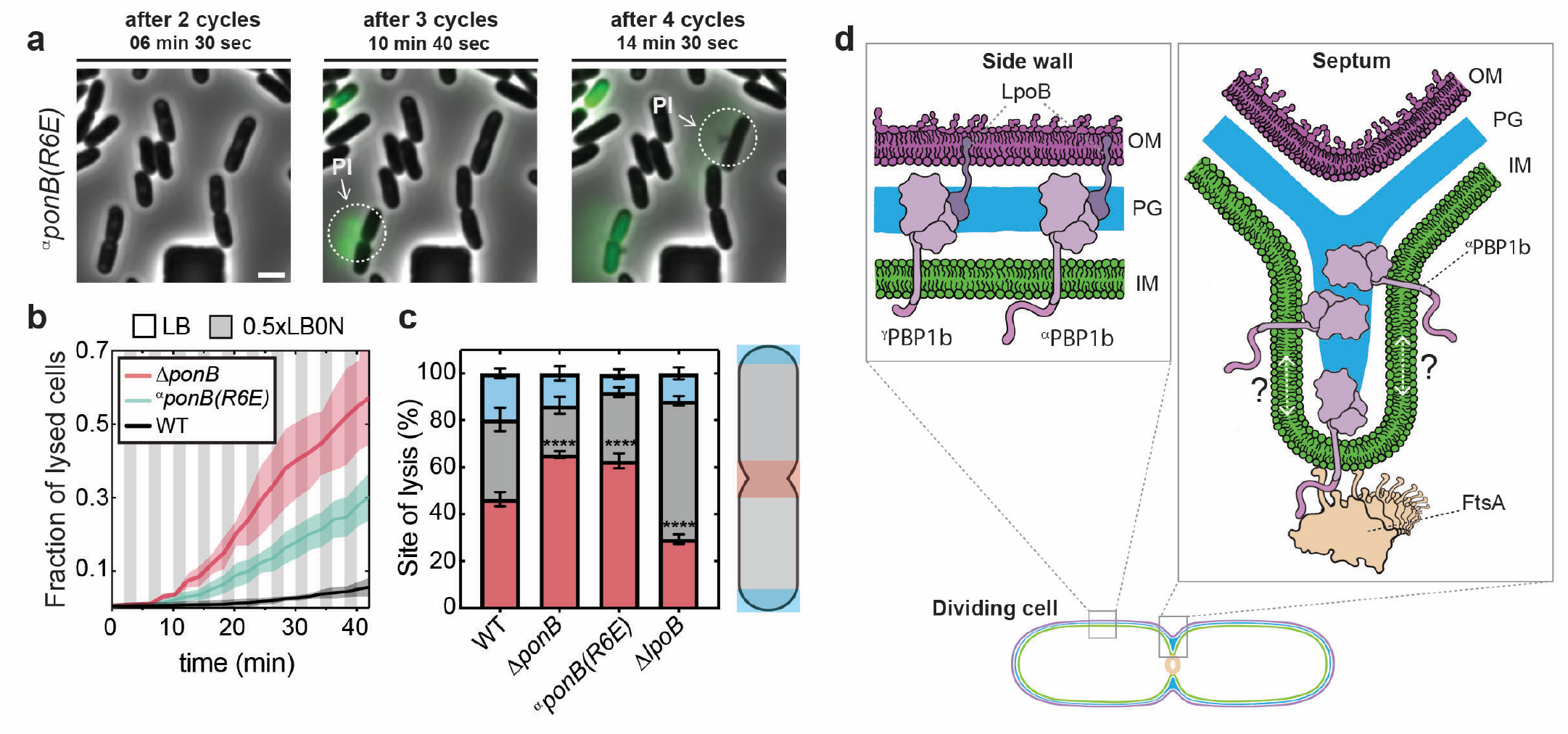
Septal localization of αPBP1b is required for its function. **(a)** Expression of the ^α^PBP1b(R6E) isoform was induced from a lactose promoter controlled construct integrated at a phage attachment site in cells harboring a deletion of the native *ponB* gene by the addition 50 µM IPTG to growth media. Representative images of cell lysis events in response to osmotic oscillations between LB and 0.5xLB0N. Visualization of lysis was aided by imaging released DNA by propidium iodide (green) staining. Arrows point towards sites of septal lysis events. Scale bar = 2 µm. **(b)** Mean fraction of lysed cells (line) ± one SD (shade) over time are plotted from N number of biological replicates: N = 3 (WT); 3 (Δ*ponB*); 5 (^α^ponB(R6E)). (c)Manual quantification of subcellular lysis sites in response to osmotic shocks from 100 cells from 3 biological replicates. Data for Δ*ponB* and Δ*lpoB* are replotted from **Fig. 3e** and is represented as mean ± one SD and significance was tested using unpaired t-test, **** = p < 0.0001. **(d)** Summary schematic showing function of PBP1b during cell elongation and division with the recruitment of ^α^PBP1b to the division site mediated by the predicted interaction with FtsA. It remains unclear where within the septum PBP1b functions (leading-edge, lagging-edge, or both) but it is presumably first recruited to the leading-edge by FtsA. Fortification of the septum by PBP1b could take place there and/or it could be released from the leading-edge to fortify regions of the septum at the lagging-edge. It is also possible that a population of FtsA molecules associates with areas of the septum other that the leading-edge to recruit PBP1b there as well.

### Conservation of the extended N-terminal peptide of αPBP1b

To assess the conservation of the ^N-pep^PBP1b amino acid sequence we searched for homologues using JackHMMER^45^ and obtained 113 hits clustered within the Enterobacteriaceae family (**Fig S9A-C**). By contrast, a similar analysis using the sequence of the glycosyl transferase (GT) domain of PBP1b resulted in 22,750 hits distributed across the entire bacterial domain. AlphaFold2 predictions indicated that selected homologs of ^N-pep^PBP1b also have the potential to interact with FtsA through the same residues as the *E. coli* peptide (**Fig. S9D**). Surprisingly, although the first 14 amino acids of these peptides are conserved between organisms, the arginine at position six was more variable than the neighboring residues P8, I9, G10, R11, suggesting that these residues also contribute to the ^N-pep^PBP1b-FtsA interaction. Importantly, non-homologous peptides from within (e.g. *Proteus* spp.) or outside (e.g. *Burkholderia, Vibrio, Shewanella* spp.) of the Enterobacteriaceae were not predicted to interact with FtsA. Notably, PBP1a homologues do not typically possess an intrinsically disordered N-terminal domain like all of the PBP1b homologues we tested, suggesting that the presence of such a domain may be predictive of septal recruitment (**Fig. S10**). Based on this sequence analysis, we conclude that division site recruitment and septum fortification is likely to be a conserved function for PBP1b among the Enterobacteriaceae with alternative recruitment mechanisms potentially promoting a similar activity in more distantly related bacteria encoding PBP1b variants with unrelated sequences at their N-termini.

## Discussion

The FtsW-FtsI complex is essential for sPG synthesis during division in most characterized bacteria^1^. By contrast, the importance of aPBPs for division varies between organisms, which has made it difficult to determine their role in the process. In Gram-positive bacteria (Firmicutes), aPBPs range from being completely dispensable for division^46–48^, to supporting sPG production along with FtsW-FtsI in others^49^, to being the essential septum synthase^8^. Similarly, in Gram-negative bacteria (Proteobacteria), interactions between *E. coli* PBP1b and divisome components^22,23^ suggest that aPBPs play important roles in the division process, but the phenotypic consequences of aPBP inactivation also varies between these organisms. There are no readily observable division defects in *E. coli* when PBP1b is inactivated whereas in *Acinetobacter baumannii* deletion of the gene encoding PBP1a results in mild cell filamentation and problems with daughter cell separation^50,51^. Even in cases where a division phenotype has been observable in cells lacking an aPBP, it is difficult to disentangle it from the general problems in PG synthesis displayed by mutants with aPBP defects. For example, PBP1b inactivation in *E. coli* results in a dramatic reduction in overall PG synthesis^14,52^. Thus, a decrease in PG labeling detected at midcell^27^ or a genetic interaction of an aPBP with a division gene^28–30^ could either reflect a direct division problem or an indirect one caused by a decline in PG content throughout the cell.

Here, we report the discovery of a division specific function for *E. coli* PBP1b. *In situ* cryo-ET indicates that this aPBP is required to produce a wedge-like structure of PG at the developing septum that contributes to its stiffness as shown by AFM. Furthermore, loss of this structure is associated with osmotic sensitivity and cell lysis via a loss of envelope integrity at the division site. Importantly, this activity of PBP1b is independent of its activator LpoB. Mutants lacking LpoB display all the general problems with PG biogenesis caused by PBP1b inactivation^10–12,16–19^ but not the defects associated with the loss of sPG wedge formation described here. Thus, our results indicate that PBP1b plays a specialized role in division site fortification that is distinct from its LpoB-dependent function in reinforcing the PG matrix throughout the cell cylinder (**Fig. 6d**). What activates PBP1b at the division site in place of LpoB is not currently known, but attractive candidates include one or more of the many PBP1b interaction partners identified previously^22–26^.

PG synthesis at the division site is thought to occur in two stages, either of which might involve sPG wedge formation by PBP1b. The first stage, referred to as pre-septal PG synthesis or PBP3(FtsI)-independent PG synthesis (PIPS), involves the localized insertion of PG at midcell to promote zonal cell elongation^53–57^. This step is followed by the FtsW-FtsI-dependent transition to the inward growth of sPG and the accompanied invagination of the membrane. Pre-septal PG synthesis occurs independently of many cell division proteins, including FtsI and FtsA, but depends critically on FtsZ and Z-ring formation^55,56^. The process also occurs in cells individually inactivated for either PBP1a or PBP1b^56^. Thus, if aPBPs indeed play a critical role in pre-septal PG synthesis as some results suggest^25^, the two different enzymes are interchangeable for this activity. By contrast, sPG wedge formation and division site stabilization were found to be a PBP1b specific function. Additionally, our results suggest that PBP1b is recruited to the division site to promote sPG wedge formation via an interaction with FtsA, which is not required for pre-septal PG synthesis^56^, and the wedge-like sPG structure was only observed in cryo-electron tomograms of cells with constricted membranes^31^. We therefore conclude that the division site stabilizing activity of PBP1b reported here is operating contemporaneously with sPG ingrowth promoted by FtsW-FtsI rather than pre-septal synthesis. How the activities of PBP1b and FtsW-FtsI are related to and/or coordinated with each other in space and time during this phase of division remains to be determined. However, an attractive possibility is that the mechanism resembles a previous proposal for Gram-positive bacteria^58^ in which FtsW-FtsI produces a core of sPG that is sandwiched by fortifying layers produced by PBP1b. Notably, it may be differences in the relative importance of the fortifying sPG layer that contributes to the variable severity of division phenotypes associated with aPBP inactivation in different bacteria.

Finally, our results not only demonstrated a division specific role for PBP1b but one that depends on a specific isoform of the enzyme (**Fig. 6d**). It has long been known that the *ponB* gene encoding PBP1b has two start codons that produce either the longer ^α^PBP1b or the shorter ^γ^PBP1b isoform^37–39^, but the functional relevance of the two forms has remained unclear. Here, we show that it is the extended N-terminus of the alpha isoform that is needed for the recruitment of PBP1b to midcell for division site fortification. Notably, a previous study found that the alpha isoform of PBP1b was specifically required to prevent cell lysis when PBP1a and FtsI(PBP3) were simultaneously targeted with beta-lactam antibiotics^39^. The authors also found that this activity involved the first six residues of ^α^PBP1b and astutely suggested that they may function to promote an interaction with the division apparatus^39^. Our results confirm this suggestion and argue that the recruitment of ^α^PBP1b to the divisome involves an interaction between ^N-pep^PBP1b and FtsA with residue R6 of ^α^PBP1b making a critical contact with FtsA. Although this interaction requires further biochemical and structural validation, the observation that high-confidence interactions are also predicted for ^N-pep^PBP1b-FtsA pairs from other enterobacteria supports the recruitment model and suggests that the mechanism is likely to be conserved. Regardless of the exact interaction partner, the extended cytoplasmic N-terminus of ^α^PBP1b is clearly required for the recruitment. Many aPBPs encoded by diverse bacteria have extended cytoplasmic N-termini, and in Firmicutes these termini are also involved in division site localization via interactions with the Gram-positive division protein GpsB^59^. Therefore, although the details are likely to differ, the division site recruitment and fortification mechanism reported here for *E. coli* PBP1b may be widespread among bacteria.

## Acknowledgements

We thank all members of the Bernhardt and Rudner labs for support and helpful conversations. We are grateful to Dr. Christel Genoud, Jean Daraspe, Antonio Mucciolo and Damien de Bellis at the Electron Microscopy Facility of the University of Lausanne and Dr. Emmanualle Jeanvoine for providing access to workstations for cryo-ET image processing. We thank Dr. Sarah Sterling, Dr. Christopher Borsa, Dr. Jenn Podgorski, Dr. Phat Vinh Dip, Dr. Edward Brignole and Dr. Anna Osherov at the MIT.nano cryo-EM facility, Dr. KangKang Song and Dr. Chen Xu at the University of Massachusetts cryo-EM facility and Dr. Richard Walsh and Dr. Zongli Li at the cryo-EM @ Harvard Medical School facility for providing access to the cryo-EM microscopes and for all their help, advice, and maintenance of cryo-EM equipment. AFM was performed at the Harvard University Center for Nanoscale Systems (CNS); a member of the National Nanotechnology Coordinated Infrastructure Network (NNCI), which is supported by the National Science Foundation under NSF award no. ECCS-2025158. We thank Dr. Nicholas S. Colella for excellent advice on AFM data acquisition and analysis. Furthermore, we would like to express our gratitude to the MicRoN imaging core at Harvard Medical School, for excellent advice on live cell imaging and maintenance of fluorescence microscopes. We are grateful to Bara Krautz for creating the cartoon illustrations (www.sciencecommunicated.com). We thank Lucy Miles for assistance with creating pLM001. A.V. was supported by a EMBO long-term postdoctoral fellowship ALTF_89-2019 and a Swiss National Science Foundation (SNSF) Post-doc.Mobility fellowship P500PB_203143. This work was also supported by funding from the National Institutes of Health (R35GM142553 to L.H.C. and R01AI083365 to T.G.B.), investigator funds from the Howard Hughes Medical Institute (T.G.B.), a SNSF Starting Grant (TMSGI3_218251 to P.P.N.) and the Foundation Pierre Mercier pour la Science (to P.P.N).

## Author contributions

P.P.N, A.V. and T.G.B. conceived the project and interpreted data. P.P.N and A.V. performed experiments, analyzed data. P.P.N. performed cryo-FIB / cryo-ET and image processing. A.V. generated mutants, performed fluorescence microscopy, AFM experiments, and bioinformatics. V.Y.A. performed 3D rendering and template matching. V.Y.A. and A.T. performed 3D segmentations of cryo-ET data. E.W.S. performed the AlphaFold interaction screen and was supervised by J.C.W. R.H. performed TIRF experiments, analyzed and interpreted data, under supervision of M.L. J.C.W., L.H.C., P.P.N., A.V. and T.G.B. provided infrastructure. T.G.B. wrote manuscript with support from P.P.N. and A.V.

## Materials and Methods

### Media, bacterial strains, and mutagenesis

All strains used in this study are derivatives of *E. coli* MG1655 are listed in **Tables S2 and S3** and primers used for PCR are listed in **Table S4**. Bacteria were grown in LB (1 % Tryptone, 0.5 % yeast extract, 0.5 % NaCl), 0.5xLB0N (0.5 % Tryptone, 0.25 % yeast extract) or M9 medium^60^ supplemented with 0.2 % D-glucose and casamino acids each as indicated. For selection, antibiotics were used at 10 µg ml^-1^ (tetracycline), 25 µg ml^-1^ (chloramphenicol), and 50 (kanamycin, ampicillin) µg ml^-1^. Mutant alleles were moved between strains using phage P1 transduction. If necessary, the antibiotic cassette was removed using FLP recombinase expressed from pCP20^61^. All mutagenesis procedures were confirmed by PCR and Sanger sequencing.

### Cryo-EM specimen preparation

Bacterial strains were grown overnight in LB media, back diluted 1:1000 and grown with shaking at 37°C, 250 rpm to an OD_600_ = 0.3. Cells were harvested by centrifugation (2 min, 5000 x g, RT) and resuspended in LB media to a final OD_600_ = 0.6. This cell suspension (3 µl) was applied to Cflat-2/1 200 mesh copper or gold grids (Electron Microscopy Sciences) that were glow discharged for 30 seconds at 15 mA. Grids were plunge-frozen in liquid ethane^62^ with a FEI Vitrobot Mark IV (Thermo Fisher Scientific) at RT, 100 % humidity with a waiting time of 13 seconds, one-side blotting time of 13 seconds and blotting force of 10. Customized parafilm sheets were used for one-side blotting. All subsequent grid handling and transfers were performed in liquid nitrogen. Grids were clipped onto cryo-FIB autogrids (Thermo Fisher Scientific).

### Cryo-FIB milling

Grids were loaded in an Aquilos 2 Cryo-FIB (Thermo Fisher Scientific). The specimen was sputter coated inside the cryo-FIB chamber with inorganic platinum, and an integrated gas injection system (GIS) was used to deposit an organometallic platinum layer to protect the specimen surface and avoid uneven thinning of cells. Cryo-FIB milling was performed on the specimen using two rectangular patterns to mill top and bottom parts of cells^63,64^, and two extra rectangular patterns were used to create micro-expansion joints to improve lamellae instability^65^. Cryo-FIB milling was performed at a nominal tilt angle of 14°-18° which translates into a milling angle of 7°-11°. Cryo-FIB milling was performed in several steps of decreasing ion beam currents ranging from 0.5 nA to 10 pA and decreasing thickness to obtain 150-250 nm lamellae.

### Cryo-electron tomography

All imaging was done on a FEI Titan Krios (Thermo Fisher Scientific) transmission electron microscope operated at 300KeV equipped with a Gatan BioQuantum K3 energy filter (20 eV zero-loss filtering) and a Gatan K3 direct electron detector. Prior to data acquisition, a full K3 gain reference was acquired, and ZLP and BioQuantum energy filter were finely tuned. The nominal magnification for data collection was of 33,000x, giving a calibrated 4K pixel size of 2.758/2.704 Å, respectively. Data collection was performed in the nanoprobe mode using the SerialEM or Thermo Scientific Tomography 6 software. The tilt range varied depending on the lamella, but generally was from -68° to 68° in 2° steps following the dose-symmetric tilt scheme. Tilt images were acquired as 8K x 11K super-resolution movies of 4-8 frames with a set dose rate of 1.5-3 e-/Å/sec. Tilt series were collected at a range of nominal defoci between -3.5 and -5.0 µm and a target total dose of 80 to 180 e^−^/Å^2^ (**Table S1**).

### Cryo-electron tomography image processing

Acquired tilted super-resolution movies were motion corrected and Fourier cropped to 4K x 5K stacks, using *framealign* from IMOD^66^. Tilt series were aligned using *etomo* in IMOD^67^ and *Dynamo*^68^. CTF-estimation was performed in IMOD. CTF-correction was performed by *ctfphaseflip* program in IMOD. CTF-corrected unbinned tomograms were reconstructed by weighted back projection and subsequently 2x, 4x and 8x binned in IMOD. Denoising of cryo-ET data was performed in cryo-CARE^70^. Bandpass filtering and summed projection of cryo-tomogram slices was performed in *Dynamo* complemented with customized MATLAB scripts^71^.

### Segmentation

Segmentation was performed on denoised tomograms using Amira (Thermo Fisher Scientific) for the cell wall signal by non-biased semi-automatic approaches. Manual annotation was required every 10 slices, then Amira’s interpolation function was applied to automatically trace slices in between. Annotation was done in 2D slices where features of interest were visible by eye. The segmented PG signal is not indicative of specific glycan strand network but rather serves as a visual guide to relevant densities assigned to the cell wall. IM and OM were segmented using MemBrain^72,73^.

### Template matching

*Dynamo*’s template matching module was used to pick ribosomes in binned 4x cryo-electron tomograms and assigned angles. The ribosome template was adjusted to the pixel size of the tomogram in *Dynamo*^71,74^. Rendering of *Dynamo* tables from template matching process were done in ArtiaX ^75^a plugging of ChimeraX^76^.

### Quantification of cryo-ET data

Summed projection images of cryo-ET tomograms were used to quantitatively measure cell dimensions at the division site^77^.

### Periplasmic space

Measurements of periplasmic space thickness were performed from center to center of opposing IM at the septum^31^. We used a customized macro in FIJI^78^ that measures thirty Euclidean distances from surface-to-surface areas^79^ in nm, e.g., from IM to IM at the septum. For these thirty single measurements the mean was calculated, yielding a final single value per septum.

### IM-OM distance

Measurements were performed in FIJI using the ‘point to point’ measuring tool. Measurements were done from IM tip to OM tip (IM-OM distance) at the septum in FIJI^78^. The difference in length between these lines was used to report division site symmetry.

### 3D rendering

3D rendering and movies were performed in ChimeraX and ArtiaX^75,76^.

### Isolation of PG sacculi for AFM

Overnight cultures of the indicated strains were back diluted 1/1000 into 50ml of fresh LB and grown at 37·C until an OD_600_ = 0.3-0.4. Cells were harvested by centrifugation (2 min, 5000 g) and fixed in ice-cold 70% EtOH for 20 min. After three washes with ddH_2_O, cells were boiled with stirring in 4% SDS for 45 min. Sacculi were further washed (3x ddH_2_O) and treated with 200 µg/ml Pronase E (Sigma Aldrich, P5147) in 10 µM Tris-pH 7.8 containing 0.5 % SDS for 2h at 60·C. Finally, sacculi were washed 3x in ddH_2_O and stored in 500µl of ddH_2_O containing 0.02% sodium azide.

### AFM

For AFM analysis, microscope slides were pretreated as previously described^80,81^. Briefly, slides were plasma cleaned (Harrick Plasma PDC-32G-2; 2 min, high setting) and coated with Vectabond^®^ (Vector Laboratories, SP-1800-7) according to the manufactures protocol. Approximately 50µl of 1/10 diluted (in ddH_2_O) sacculi were added onto a Vectabond^®^ coated slide and allowed to adhere for 5 min before rinsing 3x with ddH_2_O.

AFM was performed at the Harvard Center for Nanosacle Systems using a JPK Nanowizard IV with UltraSpeed head (Burker). Mechanical measurements were performed in QI mode with a qp-BioAC CB3 cantilever (Nano-AndMore, qp-BioAC-10) with a nominal spring constant of 0.06 N/m and 30 kHz resonant frequency. The cantilever stiffness was calibrated by measuring the thermal noise of the cantilever in ddH_2_O. First, low-resolution 10 µm overview scans of sacculi were taken using 256 × 256 pixels. For high-resolution imaging one square micrometer scans of sacculi were taken with 128 × 128 pixels, 0.2 nN set point, 1000 nm z-length, and 62.5 µm/s z-speed. Sacculi displaying a distinguishable septal band and absence of PG-folding were imaged at high-resolution (1 µm, 128 × 128 pixels).

For mechanical measurements images were analyzed in the JPK data-processing software. The effective Young’s modulus was calculated using the Hertz–Sneddon model assuming a pyramidal tip shape, a radius of 2 nm, and a Poisson ratio of 0.5. Stiffness of the division site was quantified in FIJI^78^ by detecting septal bands on low-resolution height images using default thresholding function. Two 300 × 200 nm ROIs were manually added adjacent to the septal band (**Fig. 2b-c**). Average Young’s modulus was determined and the ratio from septal band and the sidewall was taken (**Fig 2b**). Absolute stiffness values are reported in **Fig. S3**. PG pore-size was determined from high-resolution height images in FIJI^78^ (**Fig. 2d**). First images were blurred with a 1-pixel gaussian blur filter and subsequently imported into MicrobeJ^82^ for detecting pores using thresholding function. All images were rendered for publication using JPK data-processing software.

### Bacterial growth assays

To assess the effect of PBP1b inactivation on viability, cells were osmotically challenged by growth in 0.5xLB0N at 42°C. Briefly, indicated strains were grown overnight at 30°C in LB supplemented with tet10 and IPTG 250 µM when necessary. The next morning, cells were back diluted 1/1000 in fresh LB at 37·C and grown until OD_600_ = 0.3-0.4 in presence of inducer when necessary. Cells were harvested by centrifugation (2min, 5000 g) and normalized to an OD_600_ = 1.

For spot-titers, cells were plated in 10-fold dilutions on LB or 0.5xLB0N plates and incubated at 42·C for 18h. To assess minimal levels of ectopic PBP1b production needed for complementation, cells harboring the relevant P_lac_ promoter driven expression were plated on 0.5xLB0N agar containing 0.2 % glucose, 50 µM, 100 µM, or 250 µM IPTG and incubated at 42°C. Complementation was observed at 50 µM IPTG for WT PBP1b expression constructs. Growth curves in liquid medium were measured in a Tecan M-plex (Tecan) 96-well plate reader by back diluting day cultures to OD_600_ = 0.01 in LB or 0.5xLB0N. Multi-well plates were incubated with shaking at 42 °C for a total of 6 h.

### Microfluidic osmotic shock assay

Osmotic oscillations were performed to assess mechanical integrity of cell envelope similar to previously described^83,84^. Overnight cultures of the indicated strains were back diluted 1/500 into fresh LB and grown until OD_600_ = 0.3-0.4 at 37°C. Subsequently, cells were diluted to OD_600_ = 0.1 and loaded into a CellASIC bacterial microfluidic perfusion plate (Merck, B04A) according to the manufactures protocol. The plate was then transferred to a Nikon Ti2-E inverted microscope equipped with a preheated (37°C) Okolab environmental enclosure and allowed to acclimate for at least 30min. Cells were exposed to osmotic shocks by oscillating the growth medium from LB to 0.5xLB0N ten times during a 42 min observation window with a 10 s acquisition frame rate using the ONIX flow control (Merck). This resulted in a total of 10 hypo-osmotic (LB - 0.5xLB0N) and hyper-osmotic (0.5xLB0N – LB) shocks. Cell envelope integrity was monitored by adding 5µM SYTOX-Green nucleic acid stain (ThermoFisher Scientific) or propidium iodide to the growth media. Phase-contrast and fluorescence images were recorded using a Plan Apo 100x/1.45 Oil Ph3 objective lense, a Lumencor Spectra III Light Engine illumination and GFP/RFP specific filter set (Semrock quad-band dichroic LED-DA/FI/TR/Cy5/Cy7-5X-A-000 and Semrock FF01-515/30 or Semrock FF01-641/75 emission filters), a Hamamatsu ORCA-Flash4.0 V3 sCMOS camera and Nikon Elements 5.2 acquisition software. The resulting time-lapse movies were drift corrected using a customized StackReg plugin in FIJI^85,86^. Cell lysis events were quantified from counting the number of newly appearing SYTOX signals every 10^th^ frame using the ‘Find Maxima’ function in FIJI. Sites of cell lysis events were determined from phase contrast and SYTOX-Green images by three independent lab members.

### Generation of PBP1b variants

CRIM vectors^87^ for expressing N-terminal GFP/Halo fusions to ^γ^PBP1b from the P_lac_ promoter were previously generated (pHC942 and pHC949)^14^. Additionally, full-length *ponB* was amplified from genomic *E. coli* DNA using primers (ponB-alpha-BamHI and ponB-NheI_rev (**Table S4**) and cloned into pHC942 and pHC949 using BamHI (NEB, R3136T) and NheI (R3131S). To express stable ^α^PBP1b isoform the second start codon was mutated to a leucine (M46L) using primers ponB-M46L_FW and ponB-M46L_rev (**Table S4**) with Q5 Site-Directed Mutagenesis Kit (NEB, E0552S). Ultimately, this procedure resulted in the plasmid pAV13 (Halo-PBP1b-alpha) and pAV15(msfGFP-PBP1b-alpha). Similarly, the R6E point mutation was introduced for the generation of pAV35 (msfGFP-PBP1b-alpha R6E) and pLM001 (Halo-PBP1b-alpha R6E) using primers pairs ponB-alpha-R6E-pAV15_rev and ponB-alpha-R6E-pLM01_rev, respectively. For the expression of untagged PBP1b isoforms and R6E point mutants, pAV13 was digested with XbaI/PstI and re-ligated with the PCR product of ^α^ponB amplified from its backbone using primers XbaI_RBS-ponB-alpha-FW and ponB_rev_PstI (**Table S4**). CRIM vectors were transformed into a Δ*ponB* background (TU122) and integrated into the chromosome at the *att*HK022 site using the temperature sensitive helper plasmid pAH69 (ref). Single integration events were confirmed by colony PCR and the helper plasmid was cured by incubation at 37°C.

### Assessing PBP1b levels by immunoblotting

To assess expression levels for different PBP1b isoforms and point mutants, overnight cultures of the indicated strains were back diluted 1:1000 into LB and grown to OD_600_ = 0.3-0.4 at 37·C. Cells were harvested by centrifugation (2 min, 5000 g), concentrated to 3 OD_600_ units/ml and resuspended in 50µl PBS containing EDTA-free Protease Inhibitor (Roche, 11836170001) and mixed 1:1 with 2X Laemmli Sample Buffer (Bio-Rad, 1610737). Samples were sonicated and denaturated at 100·C for 10min. Total protein concentration for each sample was determined using the Non-interfering^®^ Protein Assay (G Biosciences, 786-005) according to the instructions. Five µg/ml total protein were loaded onto 4-20% Mini-PROTEAN^®^ SDS-PAGE gel (Bio-Rad, 4561094) and run at 150V for 1h. The gel was transferred to a PVDF low fluorescence membrane using the Trans-Blot Turbo Transfer System (Bio-Rad) and blocked with 5% milk in PBS containing 0.05% Tween20 (M-PBS-T). Proteins were detected with primary antibodies (polyclonal rabbit α-PBP1b serum diluted 1:10,000^10^ or monoclonal mouse α-RpoA antibody diluted 1:10,000 (BioLegend, 663104) and incubated for 1h in 1% M-PBS-T. The membrane was then washed 3 times with 0.05% PBS-T, incubated with secondary antibody solution (IRDye 800CW Anti-Rabbit IgG Goat Secondary Antibody and IRDye 680RD Goat anti-Mouse IgG Secondary Antibody, LI-COR Biosciences) in 0.5 % M-PBS-T for 1h at room temperature and washed again 3 times with 0.05% PBS-T. The blot was visualized using a fluorescence imager (BioRad, ChemiDoc MP).

### Widefield fluorescence microscopy

To visualize the localization of PBP1b isoforms, indicated strains were grown overnight in LB-tet10 at 30·C. Cells were washed 3x, back-diluted 1:1000 in M9-Glu-CAA supplemented with 50 µM IPTG and grown until OD_600_ = 0.3-0.4 at 37·C. Samples were harvested by centrifugation (2 min, 5000 g) and immobilized on a 2% agarose in M9-Glu-CAA pad and covered with a coverslip. Cells were imaged on a Nikon Ti-E inverted widefield microscope equipped with a fully motorized stage and perfect focus system and custom-made environmental enclosure heated to 37·C. Images were acquired using a 1.45 NA Plan Apo ×100 Ph3 DM objective lens with Cargille Type 37 immersion oil. Fluorescence was excited using a Lumencore SpectraX LED light engine (50ms exposure, 100% LED power) and filtered using ET-GFP (Chroma, 49002) filter set. Images were recorded on an Andor Zyla 4.2 Plus sCMOS camera (65 nm pixel size) using Nikon Elements (v5.10) acquisition software. Images were analyzed and rendered for figure or movie display with FIJI^78^ and demographs were generated using MicrobeJ (v5.13d)^82^. Original images and detection parameters were uploaded on Zenodo repository (link will be accessible upon publication).

### TIRFM for SPT of PBP1b isoforms

Overnight cultures of indicated strains were back diluted 1/1000 in LB supplemented with Tet10, Cam25 and grown at 37°C until an OD_600_ = 0.2-0.3. Janelia Fluor Halo-ligand JF549 (Promega, GA1110) was added at 20 nM final concentration, while cells were continued to incubate for an additional 20min. Subsequently, 1ml of cells was harvested by centrifugation (2 min, 5000 g) and washed 3x in 1 ml LB before resuspending in 30 µl of LB. Cells were added on a plasma cleaned (Harrick Plasma PDC-32G-2; 1 min, high setting) #1.5H coverslip (Marienfeld, 0107052) and immobilized on a 2% (w/v) agarose in LB pad.

TIRF microscopy was carried out on a Nikon Ti inverted microscope equipped with a fully motorized stage, 488 and 561 laser lines, Chroma ET GFP (49002) and mCherry (49008) filter cubes, ApoTRIF 100x 1.49NA objective with correction collar, and a stage-top incubator (Oko-lab) heated to 37°C. The microscope was controlled by Nikon Elements (4.30) and SPT movies were recorded on an Andro Zyla 4.2 Plus sCMOS camera (Oxford Instruments) using 2×2 binning with 50ms exposure and 30% laser power for a total duration of 30s resulting in an effective acquisition frame rate of 20 Hz. Single brightfield and ZapA-GFP reference image were acquired after each movie.

### SPT analysis

Particle tracking was performed in FIJI^78^ using TrackMate v7.11.1^41^. Foci were detected using the LoG detector with an estimated object diameter of 0.4 and an initial quality threshold of 10. Spurious spots outside cells were filtered using intensity-based thresholding of brightfield signal. Foci were further filtered using intensity-based thresholding of ZapA-GFP signal to compare PBP1b dynamics at the septum (high GFP signal) or the sidewall (low GFP signal). Foci were linked using the LAP Tracker with a maximum linking distance of 0.4 µm and a maximum frame gap of 2. Tracks consisting of at least 3 spots were exported and further analyzed using the Spot-ON website^43^. To estimate the populations of immobile and freely diffusing molecules, tracks were fit to a two-state model. For all conditions the fit parameters were left free, to ensure unbiased estimates. Additional SPT statistics are summarized in **Table S5**.

### AlphaFold multimer screen for PBP1b isoforms

To identify protein interaction partners for the cytoplasmic N-termini of the different PBP1b isoforms, an AlphaFold multimer screen was performed. To this end, the amino acid sequence of the PBP1b alpha (1-60: MAGNDRE-PIGRKGKPTRPVKQKVSRRRYEDDDDYDDYDDYEDEEP-MPRKGKGKGKGRKPR) and gamma (46-60: MPRKGKGKGKGRKPR) were folded with each protein in the full *E. coli* K12 reference proteome comprising 4,233 proteins (UniProt Proteome ID: UP000000625). Each pair was folded with three of the five AlphaFold-Multimer models, resulting in three independent predictions for each protein pair. Hits were defined as pairs where any of the three models had any inter-chain residues with two non-hydrogen atoms positioned within 8 Å; a predicted local distance difference test (pLDDT) > 50; and a predicted alignment error (PAE) value of < 15 Å. The results of the multimer screen are in Extended Data Table 1. This screen yielded 579 “hits” out of 8,466 pairwise interactions (442 (alpha) vs 137 (gamma), see Extended Data Table 1).

Interactions between ^N-pep^PBP1b and FtsA were further verified in selected species based on sequence conservation analysis by JackHMMER (see below) using the publicly available ColabFold v1.5.5 webserver (https://colab.research.google.com/github/sokrypton/Colab-Fold/blob/main/AlphaFold2.ipynb) using default parameters. The FtsZ-FtsA binding position (**Fig. S6**) is based on the identified site in the structure from *Thermotoga*^88^.

### Sequence and conservation analysis of PBP1b N-IDD

To assess amino acid sequence conservation of the ^N-pep^PBP1b we ran iterative HHM using JackHMMER with default parameters on the EMBL-EBI web interface (https://www.ebi.ac.uk/Tools/hmmer/search/phmmer)^89^. The search was restricted to bacteria. Obtained hits (113) were imported and displayed on using AnnoTree v1.2^90^. Randomly picked sequences of ^N-pep^PBP1b were tested for interaction with FtsA using ColabFold (see above). In contrast, running the glycosyltransferase domain of PBP1b with the same parameters results in 22750 hits distributed across the whole bacterial domain. Classical multiple sequence alignments (MSA) were carried out using Clustal Omega on the EMBL webserver (https://www.ebi.ac.uk/jdispatcher/msa/clustalo) using default parameters^91^. Predictions for intrinsic disorder were calculated using PrDOS server (https://prdos.hgc.jp/cgi-bin/top.cgi) with a 1% false positivity rate cutoff^92^. Protein structures were visualized using ChimeraX^76,93,94^. UniProt accession number of all tested proteins for *in silico* (ColabFold, MSS, IDD prediction) analysis can be found in **Table S6**.

### FtsZ treadmilling assay on supported lipid bilayers

#### Purification and Labeling of Proteins

Alexa Fluor 488-labeled FtsZ and FtsA were purified following previously established protocols^95,96^. ^N-pep^PBP1b (residues 1-64) peptide and an R6E derivative, modified with a C-terminal cysteine and an N-terminal 6x-His tag, were obtained from Biomatik. Peptide labeling was performed using a modified version of a previously described method^96^. Briefly, lyophilized peptides were dissolved in ddH2O at a concentration of 1 mg/ml and reduced with a 20-fold molar excess of TCEP for 20 minutes at room temperature. Subsequently, a thiol-reactive sulfo-cyanine5-maleimide dye (Lumiprobe), dissolved in dimethyl sulfoxide (DMSO), was added at a 5-fold molar excess and incubated for 3 hours at room temperature. Labeled peptides were then dialyzed overnight at 4°C against reaction buffer (50 mM Tris-HCl, pH 7.4; 150 mM KCl; 5 mM MgCl2). Excess unreacted dye was removed using a PD10 desalting column (Cytiva), after which peak fractions were collected, flash-frozen in liquid nitrogen, and stored at -80°C.

#### Preparation of Small Unilamellar Vesicles (SUVs)

Experiments were conducted using a lipid composition of 1,2-dioleoyl-sn-glycero-3-phospho-(1’-rac-glycerol) (DOPC) and 1,2-dioleoyl-sn-glycero-3-phospho-(1′-rac-glycerol) (DOPG) in a 2:1 molar ratio. Chloroform-dissolved lipids were combined in a glass vial, vortexed, and dried under a stream of filtered N2 to form a thin, homogeneous film. Residual chloroform was removed by placing the vials under vacuum for 3 hours. The lipid film was subsequently rehydrated in reaction buffer (50 mM Tris-HCl, pH 7.4; 150 mM KCl; 5 mM MgCl2) for 30 minutes at 37°C to achieve a final lipid concentration of 5 mM. The suspension was vigorously vortexed to form multilamellar vesicles, which were subjected to 8–10 freeze-thaw cycles using liquid N2. SUVs were generated by tip-sonicating the vesicle dispersion for 20 minutes, followed by centrifugation at 20,000 g for 5 minutes. The supernatant was collected, stored at 4°C, and used within one week.

#### Sample Preparation

Glass coverslips (#1.5H) were cleaned using piranha solution (30% H_2_O_2_ mixed with concentrated H_2_SO_4_ at a 1:3 ratio) for 1 hour, followed by extensive washing and sonication in ddH_2_O for 30 minutes. Cleaned coverslips were stored in ddH_2_O for up to one week. Prior to use, coverslips were dried with compressed air. Reaction chambers were assembled by affixing a 0.5-ml Eppendorf tube (with the conical end removed) to a coverslip using ultraviolet (UV) glue (Norland Optical Adhesive 63), followed by UV light exposure for 3 minutes.

Supported lipid bilayers (SLBs) were formed by diluting the SUV dispersion in reaction buffer to a final lipid concentration of 0.5 mM. SLB formation was induced by the addition of 5 mM CaCl_2_, followed by incubation at 37°C for 15–20 minutes. Excess non-fused vesicles were removed by washing the surface 10 times with reaction buffer. SLBs were used immediately after preparation.

#### Total Internal Reflection Microscopy (TIRFM)

All *in vitro* experiments involving SLBs were performed on a Nikon Ti2E stand equipped with an iLas2 (GATACA) 360°/Ring/Azimuthal TIRF module and a 100× NA 1.49 CFI Apochromat oil immersion objective. Fluorophores were excited using 488 nm and 640 nm laser lines from an Omicron LightHUB Ultra laser system. Emitted fluorescence was split using an TwinCam Cube 643nm and further filtered with a 525/50 ET Bandpass and a 635 nm Longpass-Filter. Time series were recorded using Andor iXon Life 888 Back-Illuminated EMCCD cameras.

#### Peptide-Filament Interaction Experiments

Self-organization of FtsA and FtsZ into treadmilling filament networks on SLBs was performed using FtsA (0.4 µM) and Alexa Fluor 488-labeled FtsZ (1.25 µM) in 100 µl of reaction buffer supplemented with 4 mM ATP and 4 mM GTP. To minimize photobleaching during imaging, the reaction buffer was supplemented with 30 mM D-glucose, 0.050 mg/ml glucose oxidase, 0.016 mg/ml catalase, 1 mM dithiothreitol (DTT), and 1 mM Trolox.

To investigate the effect of ^N-pep^PBP1b peptide on the treadmilling filament network, FtsA and FtsZ were incubated until large-scale filamentous patterns emerged. The dynamic filament pattern was monitored for 15 minutes, with image acquisition at one frame per 2 seconds and an exposure time of 50 ms for both the 488 nm and 640 nm channels. During imaging, either ^N-pep^PBP1b WT or ^N-pep^PBP1b (R6E) variants (both labeled with cyanine 5, 0.4 µM) were introduced into the reaction chamber.

#### Image analysis

For data analysis, movies were imported into FIJI (ImageJ). All micrographs presented in this study were processed using the walking average plugin in ImageJ, averaging the signal of two consecutive frames to enhance visual clarity while preserving relative intensities over time. LUTs of PBP1b peptide micrographs were adjusted in Adobe Illustrator to ensure color consistency. For intensity-over-time projections in **Figure 5d**, time-lapse frames captured during sample addition were removed. The resulting data were then normalized to the maximum intensity value for individual replicates. For colocalization data analysis Pearson’s correlation coefficient (PCC) analysis was performed using numpy.corrcoef^97^ (**Fig. S7**).

## Statistical analysis

All data measurements were plotted and analyzed using GraphPad Prism 10 (Version 10.2.2). In general, (log-) normal distribution was tested by using Shapiro-Wilk test, for comparisons of two groups, significance was determined by two-tailed, unpaired Student’s t test with Welch correction and one-way ANOVA test was used for comparison of more than two groups using the recommended post-test for selected pairwise comparisons. P values less than 0.05 were considered statistically significant. Further details on statistical tests such as sample size and number of biological repeats are provided in figure legends. For colocalization data analysis, Pearson’s correlation coefficient (PCC) analysis was performed using numpy.corrcoef^97^ (**Fig. S7**).

## Data Availability

The data, plasmids and strains that support the findings of this study are available from the corresponding authors by request. Representative tomograms are deposited in EMDB: EMD-27479 (wild-type), EMD-XYZ (Δ*ponB*), EMD-XYZ (Δ*lpoB*) and EMD-XYZ (Δ*ponA*). Corresponding raw movie frames and stacks of tilt-series are deposited as EMPIAR-11090 (wild-type), EMPIAR-XYZ (Δ*ponB*), EMPIAR-XYZ (Δ*lpoB*) and EMPIAR-XYZ (Δ*ponA*) and will be released upon publication.

## Code Availability

Scripts used in this study were deposited on Github: https://github.com/NavarroVettiger/Navarro-et-al_2022 and https://github.com/virlya-nanda/EM-ImageProcessing.

## Supplemental material

**Figure S1:**
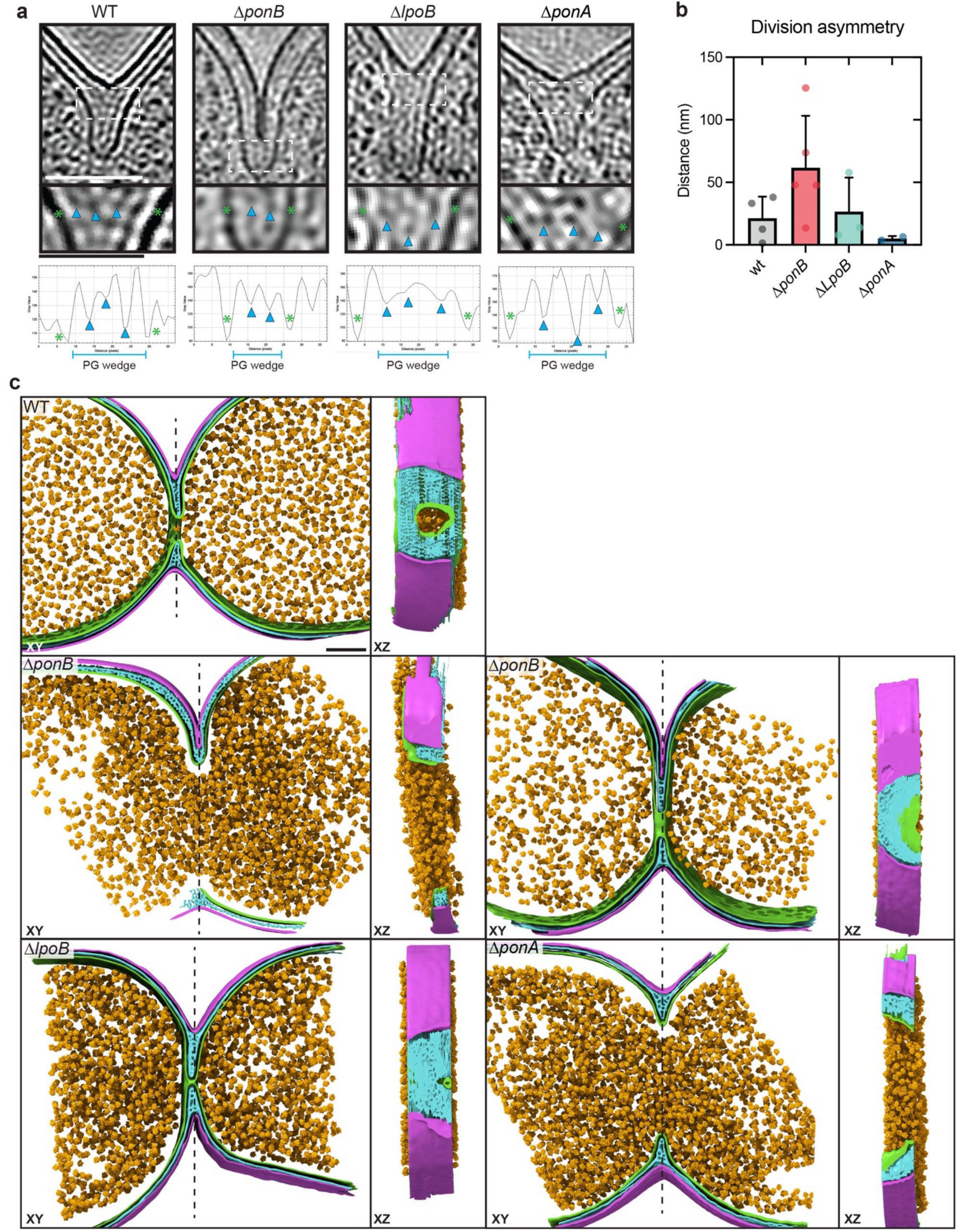
Detailed architecture of the division site by *in situ* cryo-ET. **(a)** Summed, projected central slices of low-passed filtered cryo-electron tomograms shown in Figure 1. White dashed box indicates the region of corresponding zoom-in images of the typical sPG wedge-containing region below. Bottom row shows corresponding normalized grey-scale profiles of the sPG wedge region. Green asterisks indicate IM regions and blue arrows indicate PG. Note the reduced number of detectable PG densities within the septum of the Δ*ponB* cell. **(b)** Bar graph showing the difference in nm between the length from OM to IM of both sides of the division. This denotes division side symmetry. **(c)** 3D surface segmentation renderings of IM (green), PG (cyan), OM (magenta) and ribosomes (yellow) are shown as top view (XY plane) and side view (XZ plane). Dashed line indicates the surface cut made to show a cross view of the PG architecture at the division site in the corresponding side views (XZ). N values for each strain are: N = 5 (WT); 5 (Δ*ponB*); 3 (Δ*lpoB*) and 2 (Δ*ponA*). Scale bars = 100 nm.

**Figure S2:**
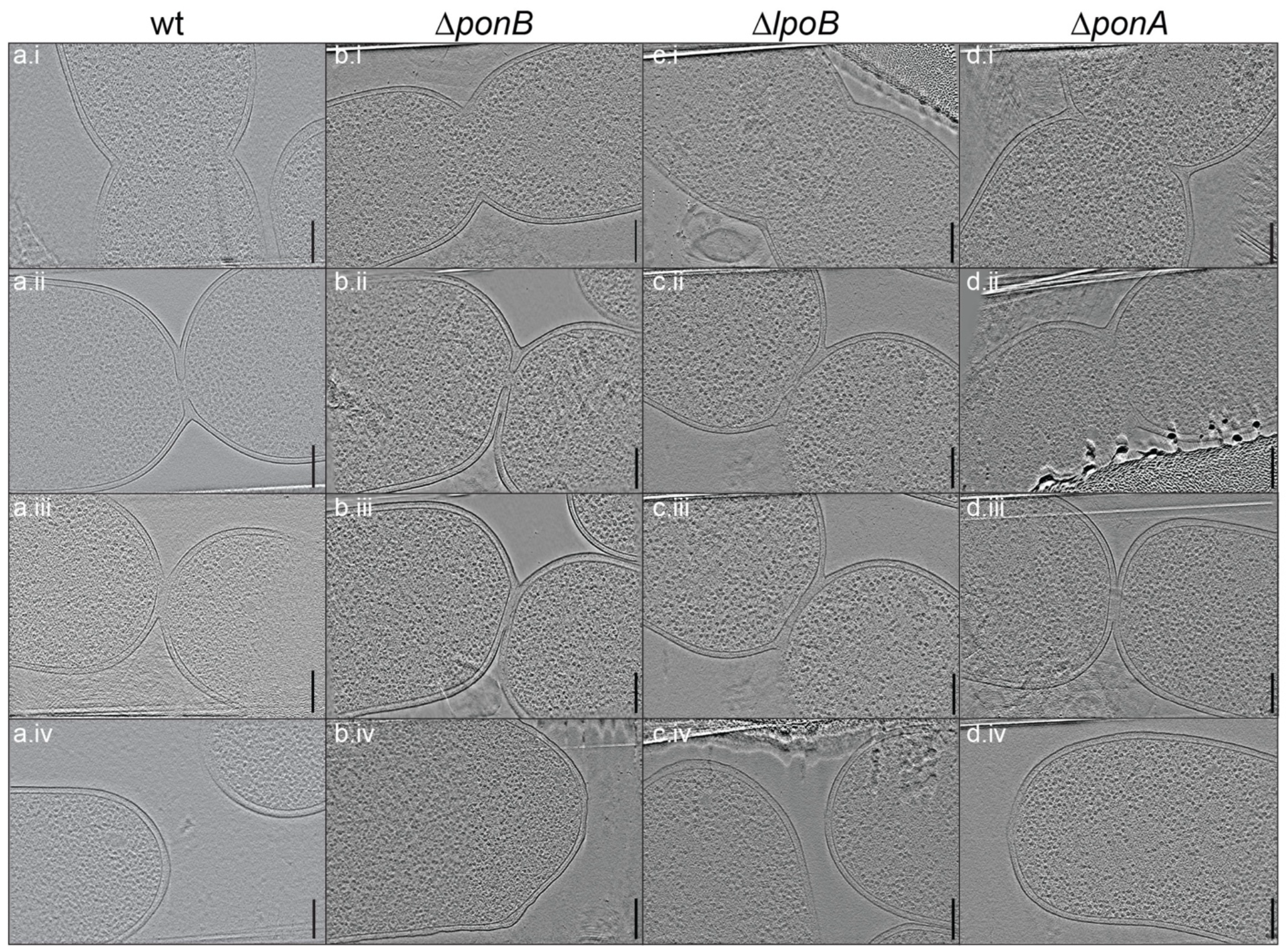
Gallery of cryo-ET data of dividing *E. coli* cells. Three dimensional slices visualizing the division site and pole of the indicated strains. Scale bar = 200 nm. A complete overview of number of tomograms and data acquisition is reported in **Table S1**. WT data from [31]. Scale bars = 200 nm.

**Figure S3:**
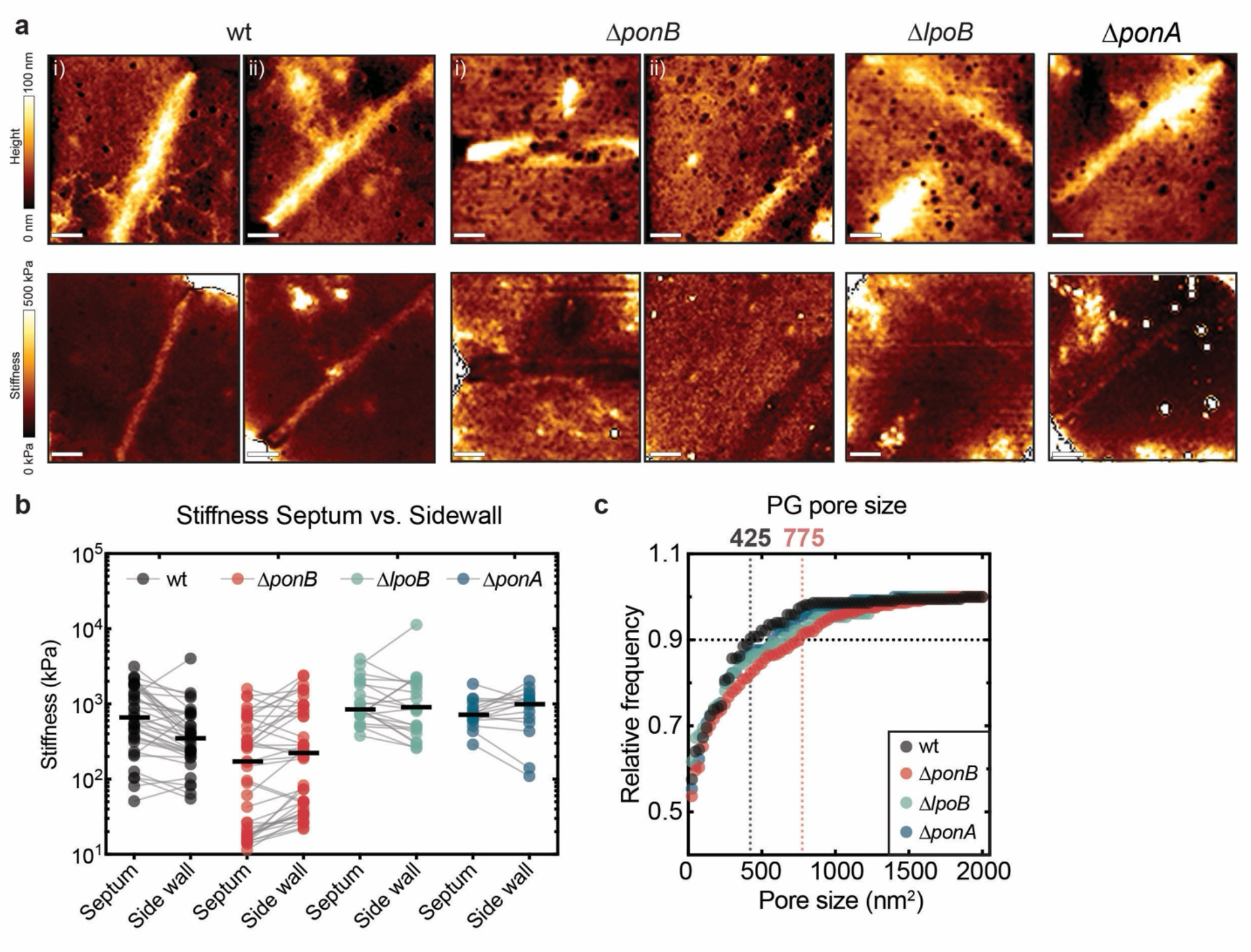
Septal ultrastructure and stiffness characterization of isolated PG sacculi by AFM. **(a)** Additional examples of high-resolution (1 × 1 µm, 7.81nm pixel size) height (top) and Young’s modulus (stiffness, bottom) maps of indicated strains. Scale bar = 200 nm. **(b)** Absolute quantification of septal and sidewall stiffness of indicated strains. Lines connect measurements from the same sacculi. N sacculi for each strain from 3 biological replicates were measured: N = 37 (WT); 40 (Δ*ponB*); 21 (Δ*lpoB*); 15 (Δ*ponA*). **(c)** Relative frequency of PG pore size as determined from high-resolution height images (see Methods). Top number displays average pore size of the top 90% of identified pores, indicating that cells lacking PBP1b display larger pores in addition to having more of them. Numbers of pores measured for each strain were: 254 (WT); 508 (Δ*ponB*); 144 (Δ*lpoB*); 147 (Δ*ponA*).

**Figure S4:**
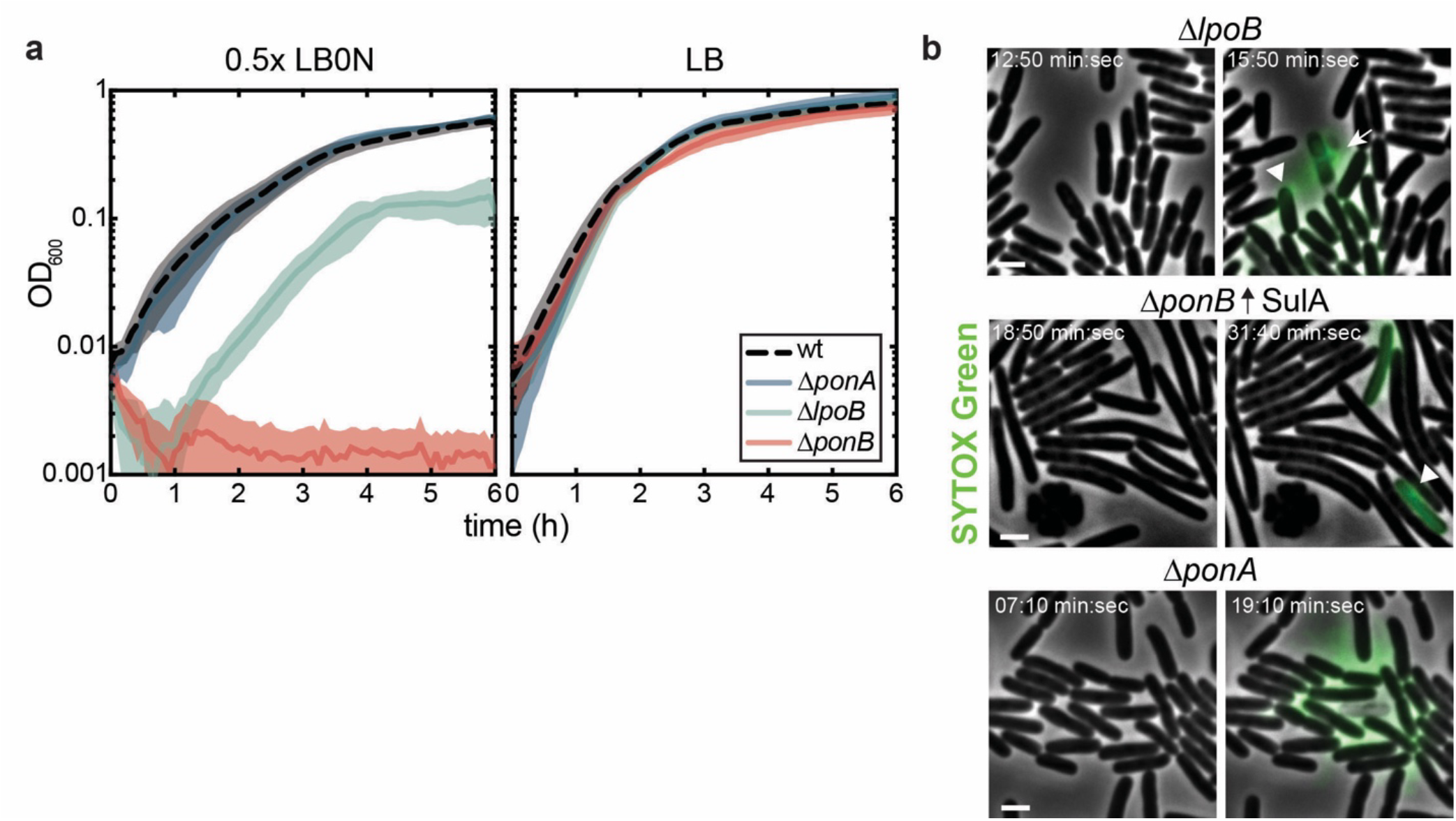
Cells lacking PBP1b are sensitive to osmotic shifts. **(a)** Bacterial growth was assessed in indicated strains using on a plate reader at 42°C in LB or 0.5xLB0N as indicated. Line represents mean ± one SD absorbance value obtained from three biological replicates. **(b)** Representative images of additional mutants imaged in response to osmotic oscillations (see Fig. 3b). Filamentation was induced by expression *sulA* from pNP146 for 20 min prior exposure to first osmotic shock. White arrows point to sites of septal lysis events, while arrowheads display polar or sidewall lysis events.

**Figure S5:**
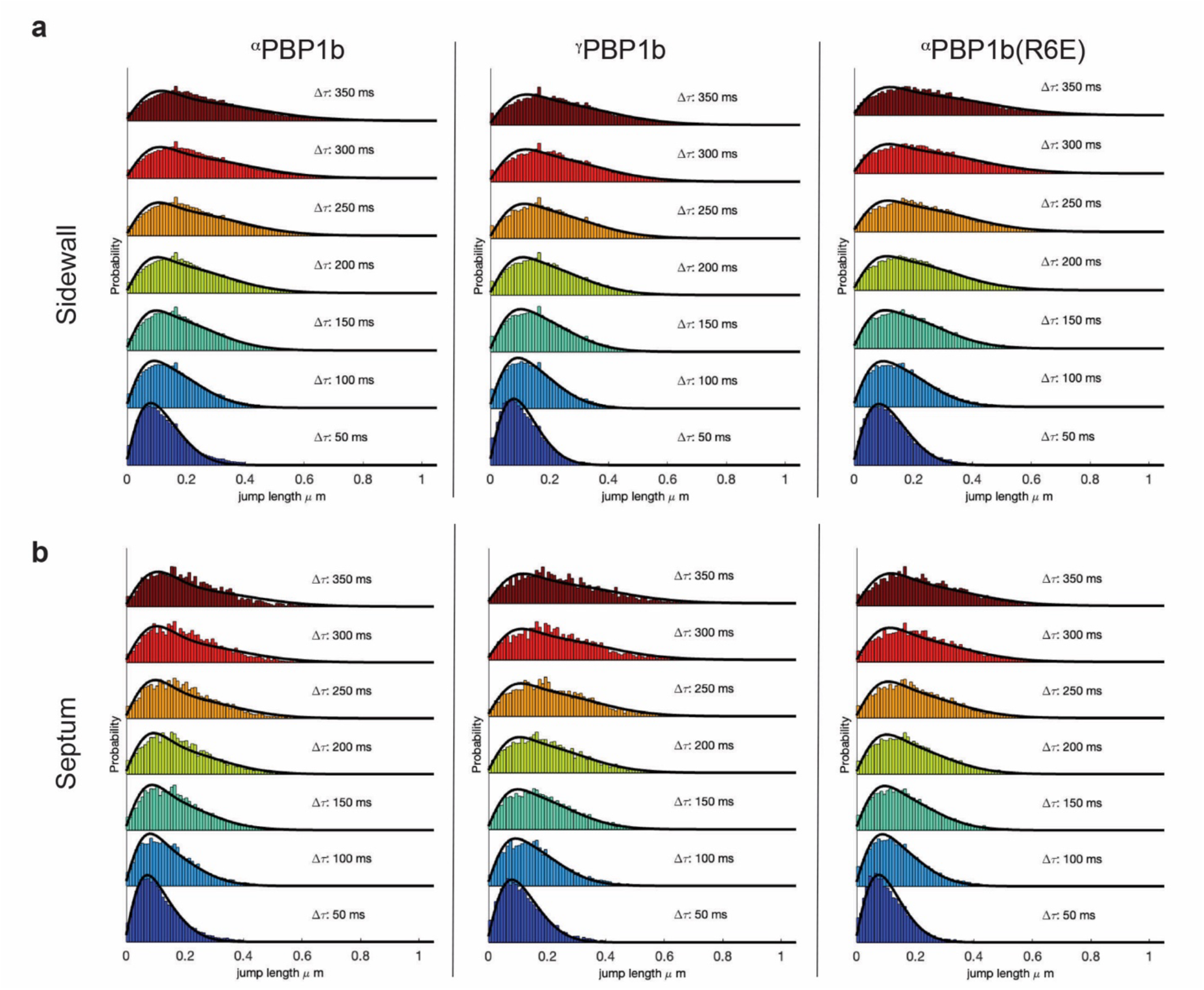
Quantification of the stationary and diffusive fraction of single particle trajectories. Observed and fit distributions of particle jump lengths at the **(a)** side wall or **(b)** septum over eight steps with a 20 Hz acquisition frame rate (Δt = 50ms) each, obtained using the Spot-On tool^43^.

**Figure S6:**
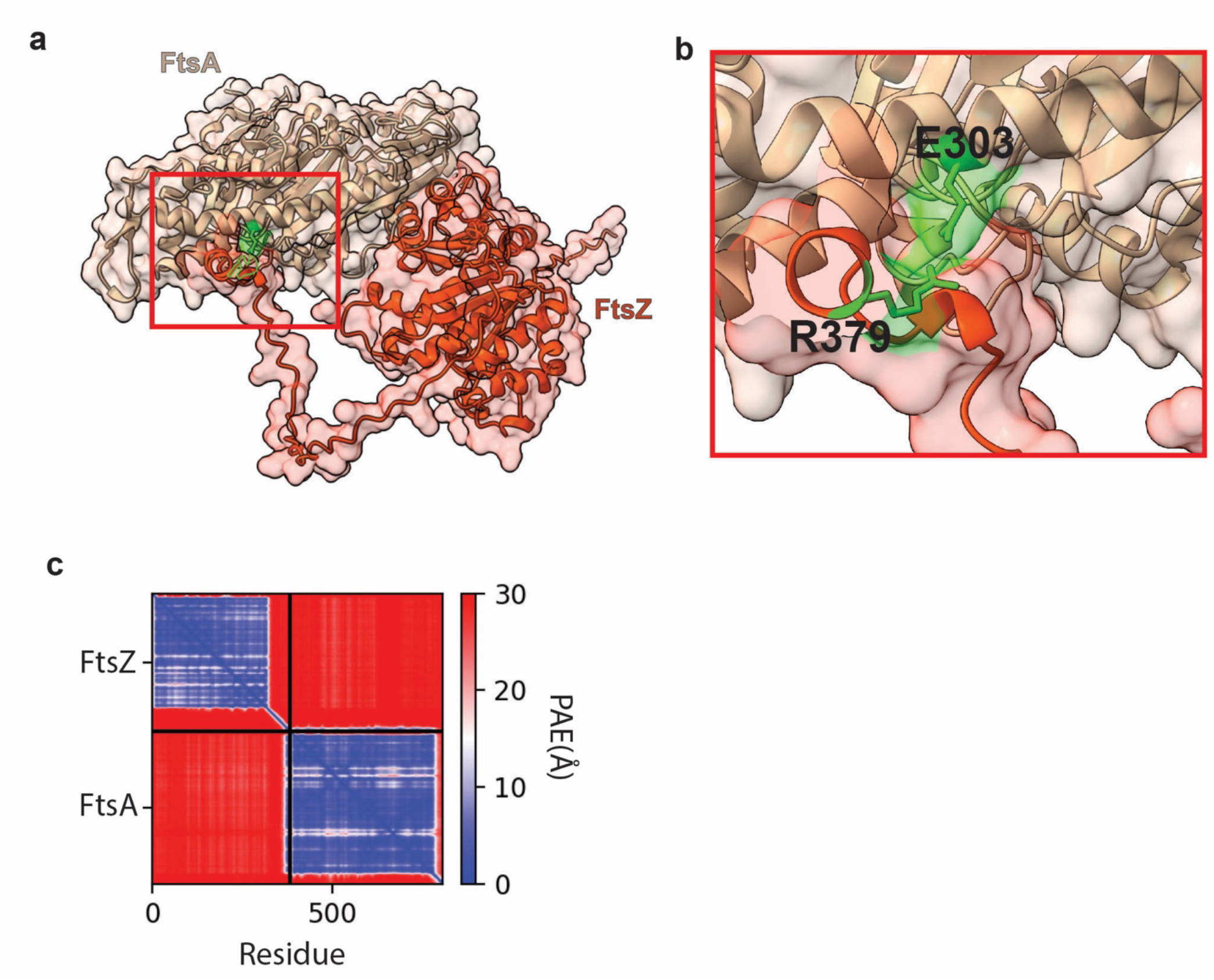
The FtsZ-FtsA interaction interface overlaps with the predicted ^N-pep^PBP1b-FtsA interface. **(a)** Structural model of FtsA (salmon) and FtsZ (red) interaction. Red box highlights the magnified region shown in b. **(b)** Magnified image of **(a)** showing the interaction between FtsA E303 residue and FtsZ R379 residue (green). **(c)** Predicted alignment error in alignment error in Å of all residues against all residues. Low error (blue) corresponds to well-defined relative domain positions. The binding position identified in the model mirrors that identified in the *Thermotoga* structure^88^.

**Figure S7:**
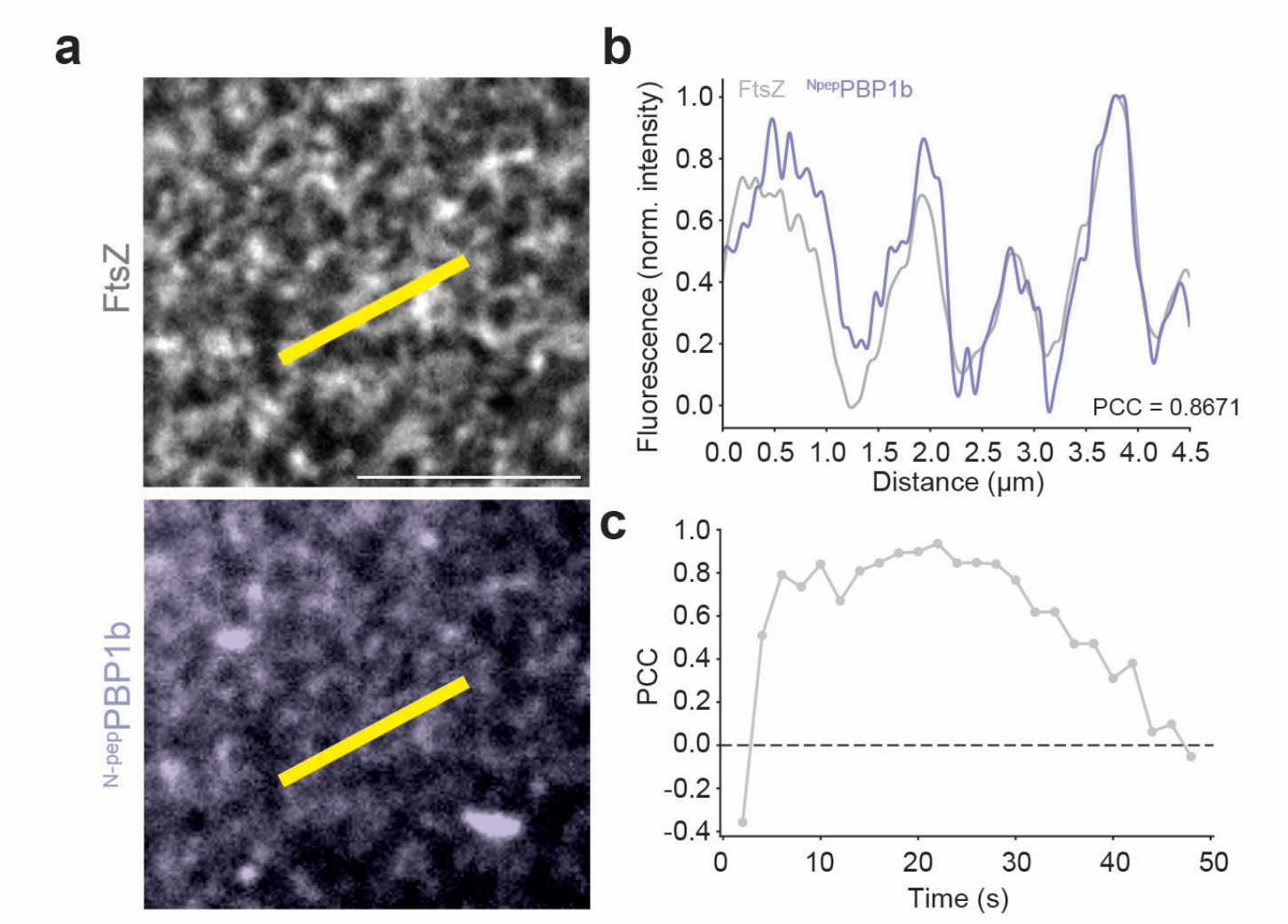
Transient colocalization of ^N-pep^PBP1b and FtsZ filaments. **(a)** Micrograph of single frame 10 seconds after addition of ^N-pep^PBP1b. **(b)** Fluorescent intensity profiles from AF488-FtsZ and Cy5-^N-pep^PBP1b corresponding to line ROI indicated in panel a. PCC was performed using numpy.corrcoef()^97^using the standard 2×2 correlation matrix of interpolated intensity profiles. **(c)** PCC values over time between normalized intensities (maximum intensity normalization) of AF488-FtsZ and Cy5-^N-pep^PBP1b in ROI from panel (a), t=0 s marks addition of Cy5-^N-pep^PBP1b.

**Figure S8:**
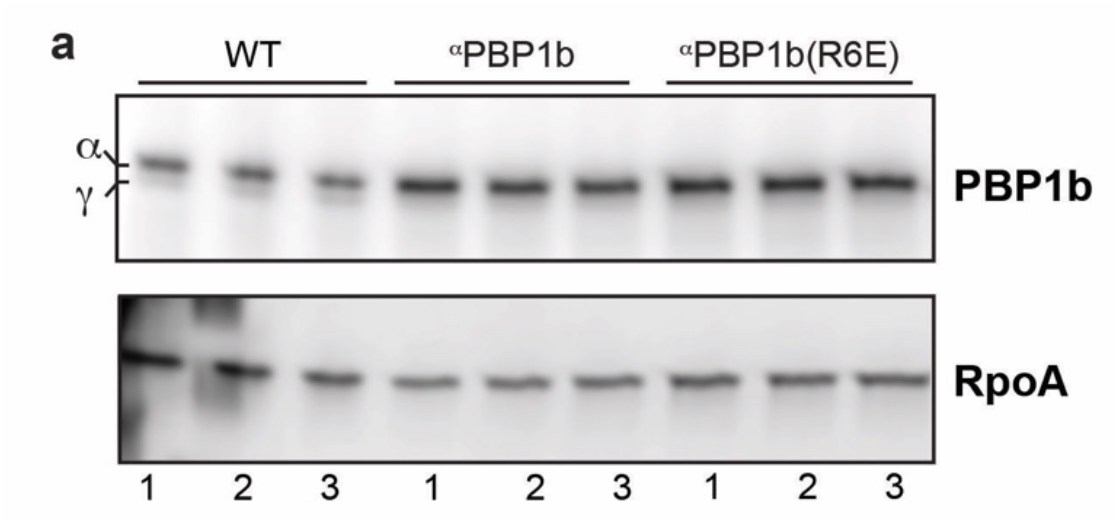
PBP1b isoforms are expressed at similar levels to WT. **(a)** Protein levels were assessed by Western blot using polyclonal rabbit anti-PBP1b serum^10^. Samples were from WT cells expressing native PBP1b and Δ*ponB* cells expressing the indicated ^α^PBP1b variant at the same induction levels used for complementation experiments. RpoA served as a loading control. Molecular weight for ^α^PBP1b is 94.2 kDa (upper band) and for ^γ^PBP1b 88.9 kDa (lower band), respectively. The molecular weight of RopA is 36.5 kDa. Protein sample from three biological replicated are displayed.

**Figure S9:**
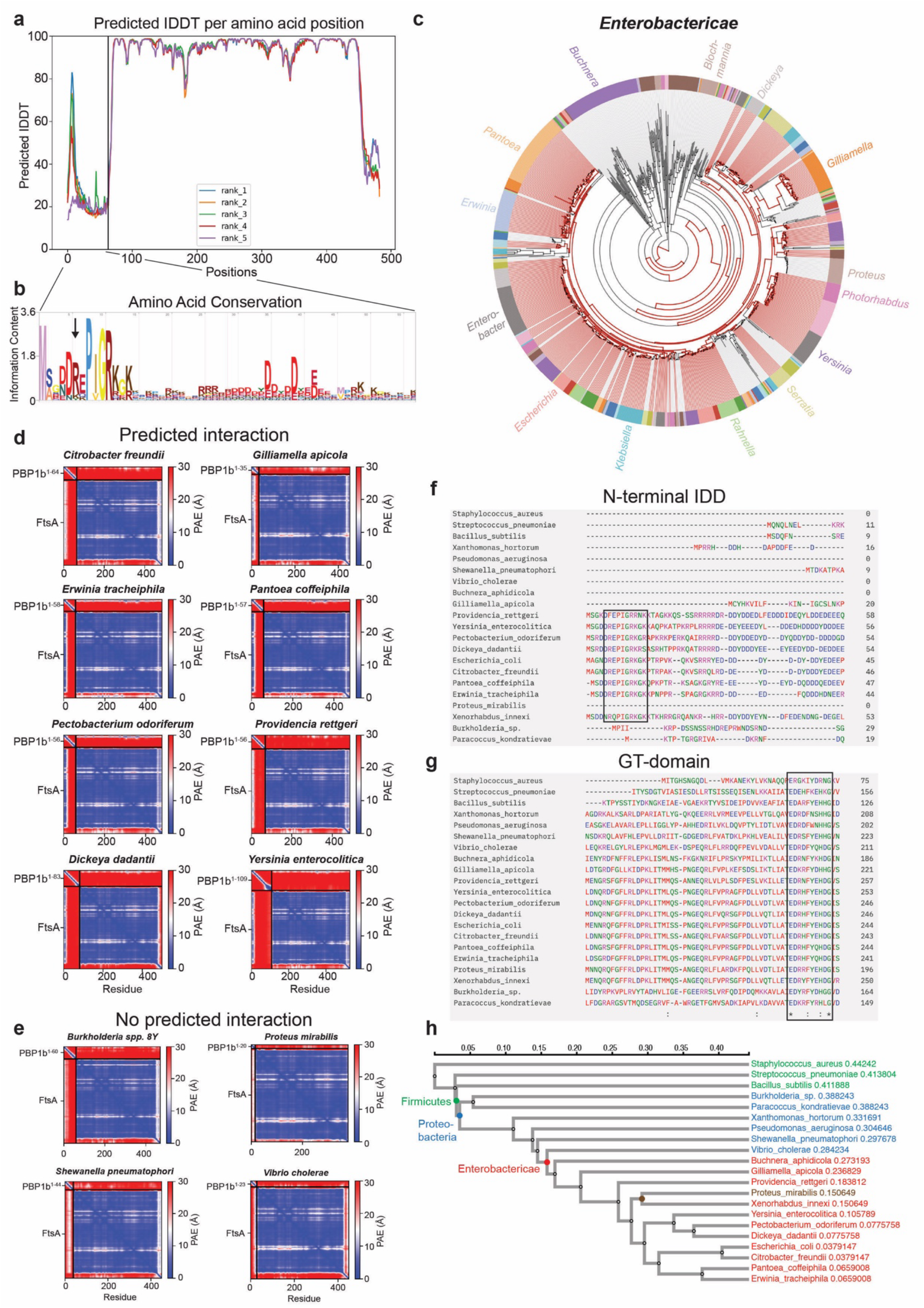
Conservation analysis of ^N-pep^PBP1b and its potential interaction with FtsA. **(a)** Predicted local distance difference test of ^N-pep^PBP1b and FtsA. **(b)** Amino acid conservation of ^N-pep^PBP1b sequence as identified by JackHMMER. Black arrow indicates R6. **(c)** Hits (in red) from JackHMMER^89^ search were visualized on a tree displaying the Enterobacteriaceae family using AnnoTree v1.2^90^. **(d)** Additional examples of high-confidence predicted ^N-pep^PBP1b-FtsA complexes in distantly related Enterobacteriaceae. **(e)** AlphaFold predictions of ^N-pep^PBP1b-FtsA complexes using ^N-pep^PBP1b from Enterobacteriaceae family members not identified in the JackHMMER search as homologues (e.g. *Proteus mirabilis*) or PBP1b proteins found in other proteobacteria outside the Enterobacteriaceae family. Multiple sequence alignment (MSA) of full-length PBP1b proteins show amino acid conservation among distantly related bacteria for the **(f)** ^N-pep^PBP1b and the **(g)** catalytic GT domain. Black box highlights conserved amino acid sequence in a (sub)set of samples. **(h)** Phylogenetic tree reconstructed for MSA. Note the absence of ^N-pep^PBP1b sequence conservation in *P. mirabilis* (brown) (see also MSA in f).

**Figure S10:**
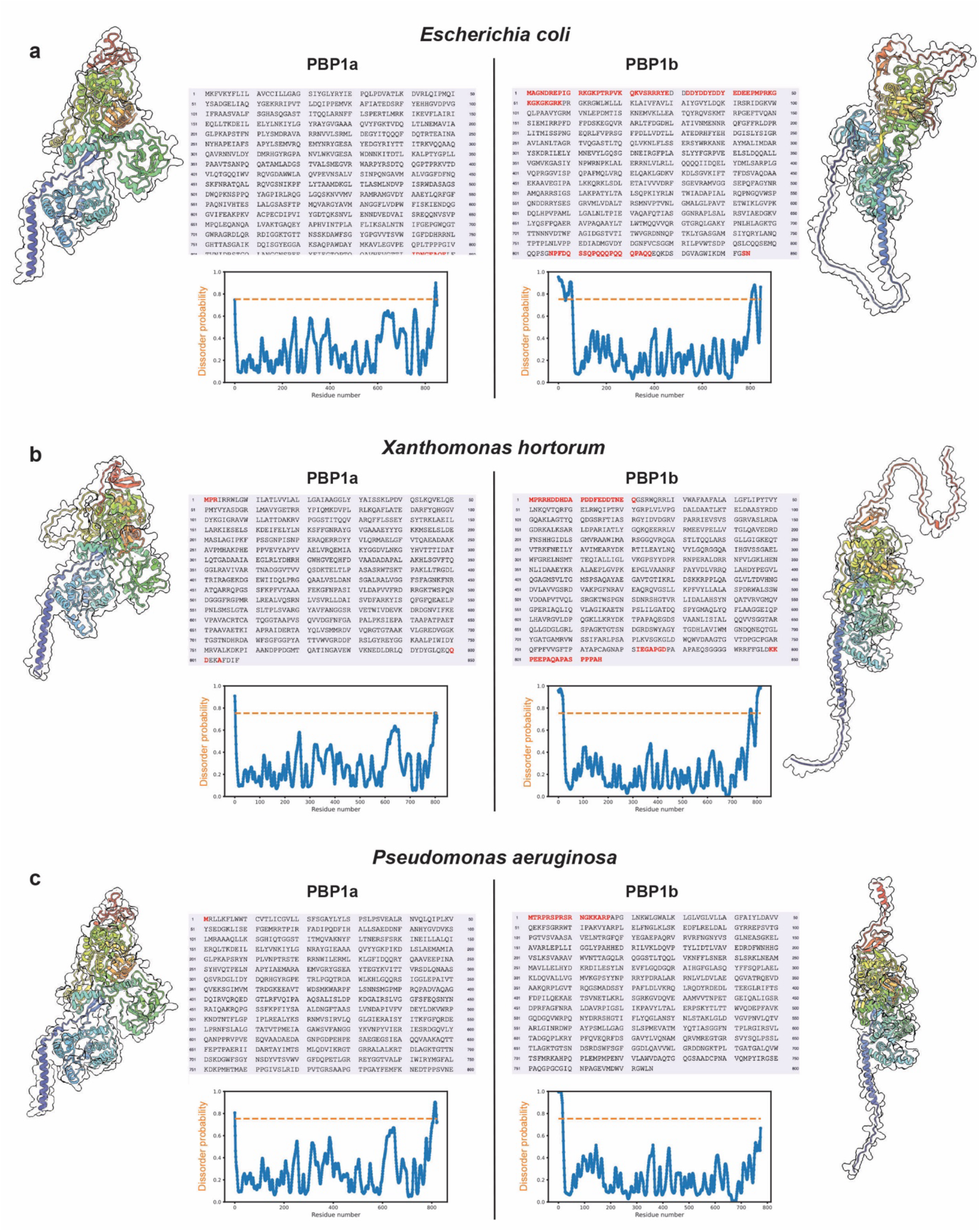
IDD domain prediction on aPBPs. AlphaFold and PrDOS highlights the presence of an on average 20-60 amino acid long N-terminal intrinsically disordered domain on PBP1b homologues which is absent on PBP1a homologues. PrDOS output displays predicted (< 1% false positivity cutoff) disordered domains in red on the amino acid sequence. Protein structures were represented in ChimeraX and are rainbow colored depending on amino acid position from their N-terminus (blue) to C-terminus (blue). Representative images for (a) *E. coli* (Enterobacteriaceae), (b) *Xanthomonas hortorum* (Enterobacterales), and (c) *Pseudomonas aeruginosa* (Pseudomonata) are displayed.

**Table S1.**
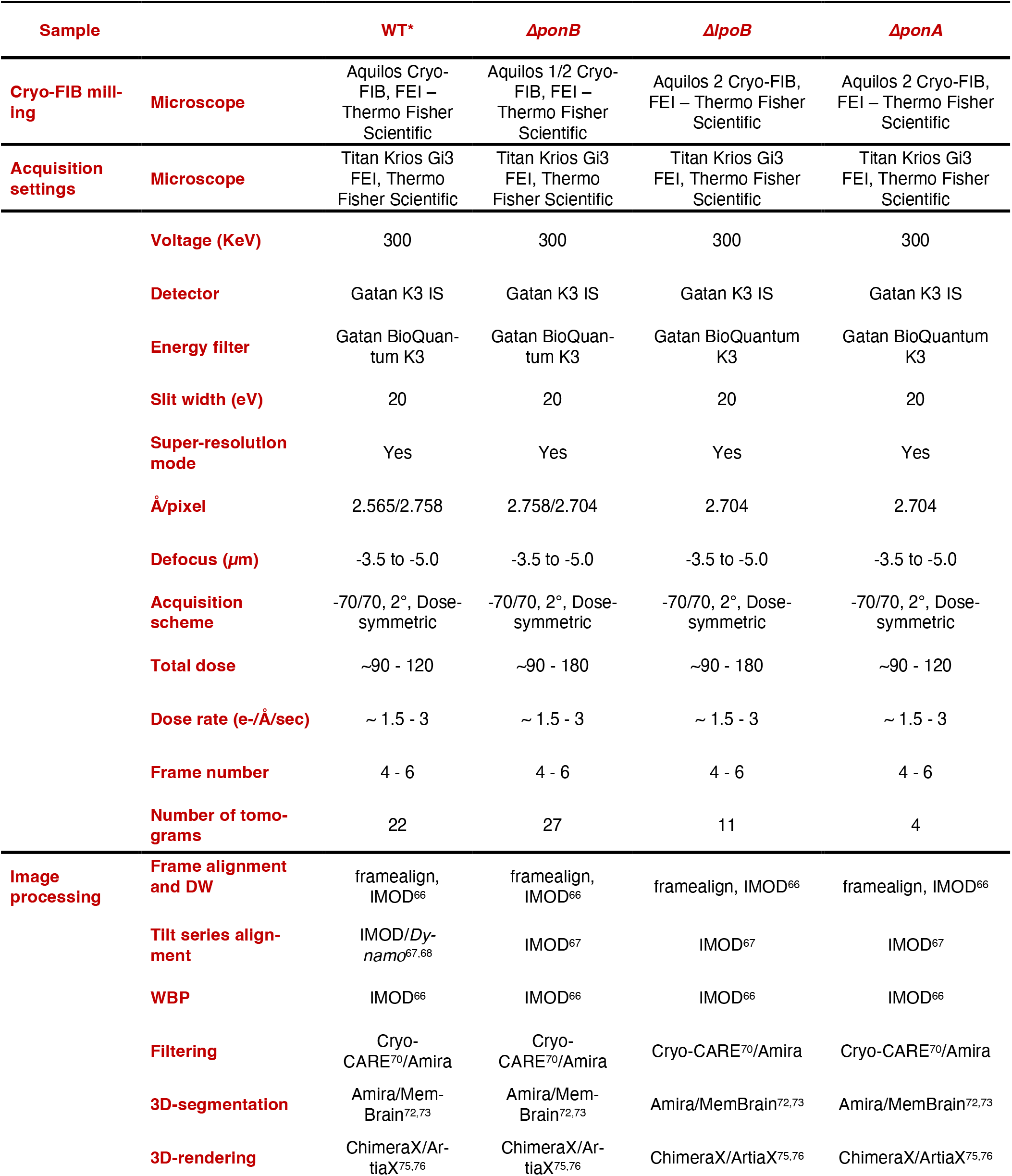
Summary of data acquisition and image processing for cryo-ET data in this study.

**Table S2.** Strains used in this study.

| Strain | Genotype | Source/Reference |
| --- | --- | --- |
| TB28 | *rph1 ilvG rfb-50 ΔlacIZYA<>frt* | [98] |
| TU121 | *TB28 ΔponA<>frt* | [10] |
| TU122 | *TB28 ΔponB<>frt* | [10] |
| CB26 | *TB28 ΔlpoB<>frt* | [10] |
| AV258(attHKAV13) | *TU122 (P<sub>lac</sub>::halo-αponB(M46L))* | P1 (attHKAV13) x TU122 |
| AV264(attHKHC942) | *TU122 (P<sub>lac</sub>::msfgfp-γponB(46-844))* | P1 (attHKHC942)<sup>14</sup> x TU122 |
| AV265(attHKHC949) | *TU122 (P<sub>lac</sub>::halo-γponB(46-844))* | P1 (attHKHC949)<sup>14</sup> x TU122 |
| AV266(attHKAV15) | *TU122 (P<sub>lac</sub>::msfgfp-αponB(M46L))* | P1 (attHKAV15) x TU122 |
| AV269 | *TB28 ΔponB<>frt, pNP146* |  |
| AV276(attHKAV13) | *AV258 zapA-sfgfp cat* | P1 (HC261)<sup>99</sup> x AV258 |
| AV276(attHKHC949) | *AV265 zapA-sfgfp cat* | P1 (HC261)<sup>99</sup> x AV265 |
| AV347(attHKAV35) | *TU122, (P<sub>lac</sub>::msfgfp-αponB(R6E, M46L))* | P1 (attHKAV335) x TU122 |
| AV352(attHKAV39) | *TU122, (P<sub>lac</sub>::αponB(R6E, M46L))* | P1 (attHKAV39) x TU122 |
| AV406(attHKLM01) | *TU122, (P<sub>lac</sub>::halo-αponB(R6E, M46L))* | P1 (attHKLM01) x TU122 |

**Table S3.**
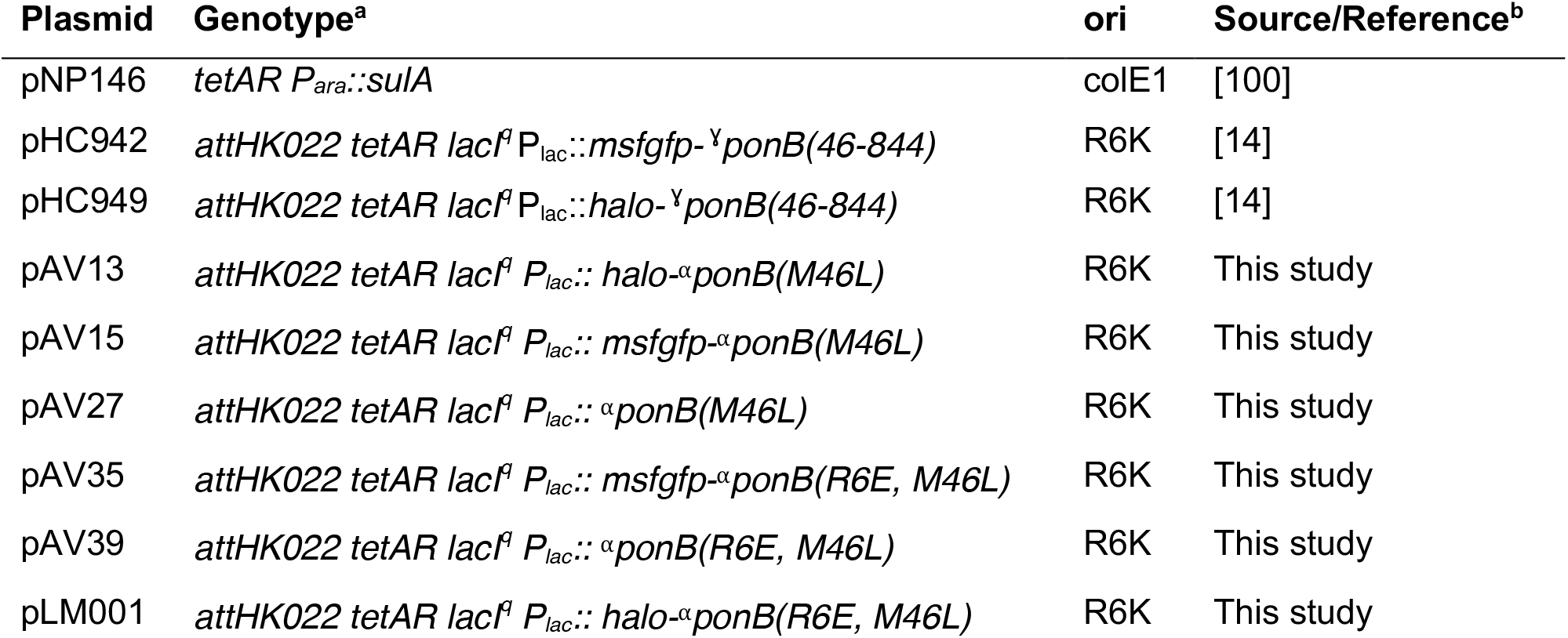
Plasmids used in this study.

**Table S4.**
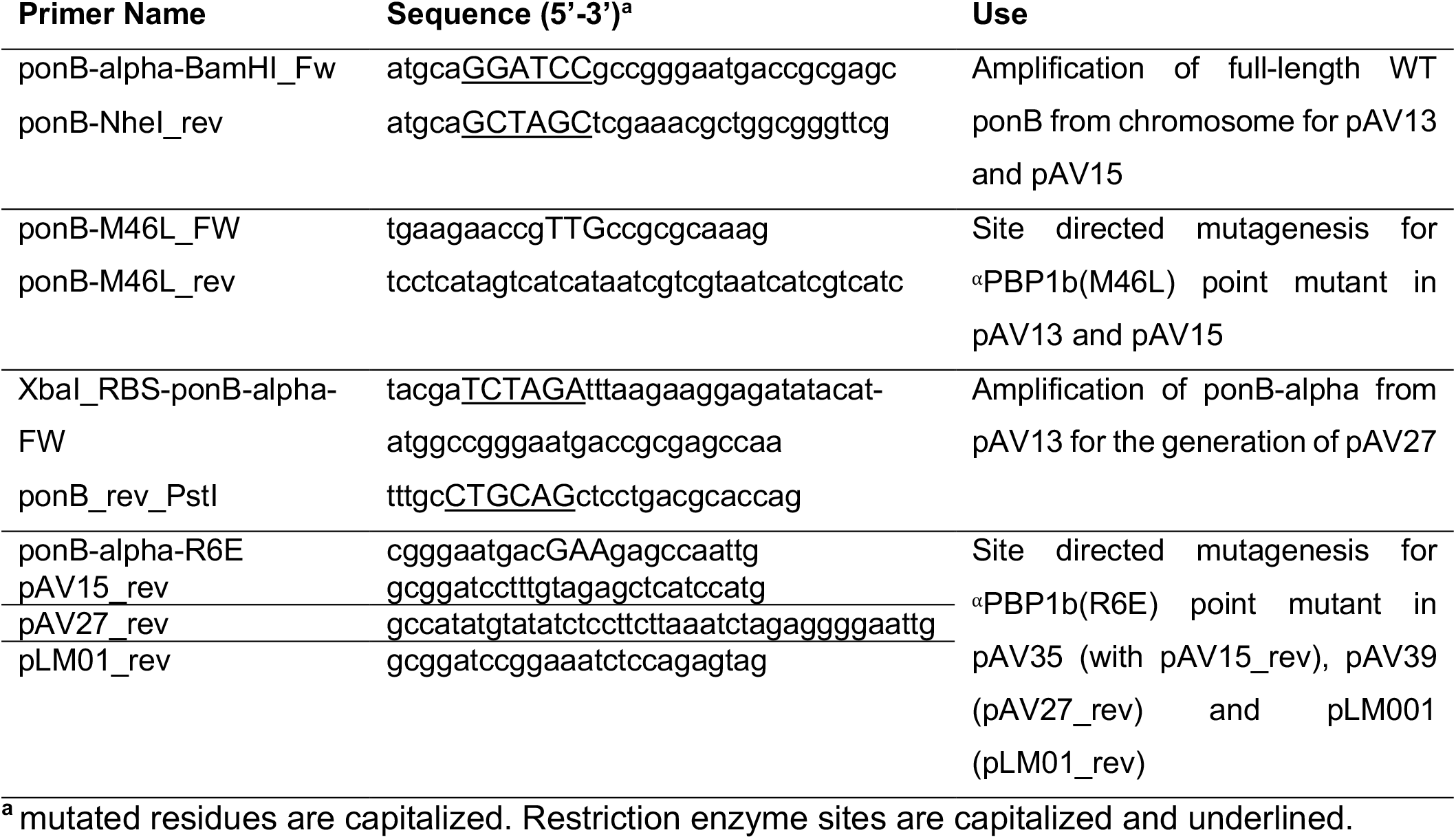
Primers used in this study.

**Table S5.** SPT statistics obtained from SpotOn^43^.

| Construct | Condition | Total trajectories<sup>a</sup> | Bound fraction | D<sub>Free</sub> (μm<sup>2</sup> sec<sup>-1</sup>) |
| --- | --- | --- | --- | --- |
| Halo-<sup>α</sup>PBP1b | Septum | 349 | 0.227 | 0.081 |
|  |  | 485 | 0.223 | 0.084 |
|  |  | 575 | 0.252 | 0.079 |
|  |  | 461 | 0.227 | 0.075 |
|  |  | 599 | 0.224 | 0.084 |
|  |  | 1029 | 0.215 | 0.081 |
|  | Side Wall | 2873 | 0.150 | 0.092 |
|  |  | 8060 | 0.153 | 0.087 |
|  |  | 5331 | 0.164 | 0.088 |
|  |  | 4531 | 0.138 | 0.089 |
|  |  | 5994 | 0.134 | 0.099 |
|  |  | 8957 | 0.155 | 0.094 |
| Halo-<sup>γ</sup>PBP1b | Septum | 981 | 0.183 | 0.086 |
|  |  | 445 | 0.115 | 0.065 |
|  |  | 699 | 0.172 | 0.073 |
|  |  | 559 | 0.112 | 0.075 |
|  |  | 676 | 0.137 | 0.076 |
|  |  | 860 | 0.173 | 0.071 |
|  | Side Wall | 6716 | 0.138 | 0.073 |
|  |  | 2695 | 0.139 | 0.064 |
|  |  | 3027 | 0.118 | 0.067 |
|  |  | 4052 | 0.155 | 0.074 |
|  |  | 3158 | 0.128 | 0.080 |
|  |  | 3555 | 0.097 | 0.079 |
| Halo-<sup>α</sup>PBP1b(R6E) | Septum | 1031 | 0.114 | 0.085 |
|  |  | 314 | 0.112 | 0.083 |
|  |  | 388 | 0.202 | 0.087 |
|  |  | 419 | 0.186 | 0.080 |
|  |  | 996 | 0.130 | 0.087 |
|  |  | 652 | 0.186 | 0.084 |
|  | Side Wall | 6190 | 0.121 | 0.094 |
|  |  | 1325 | 0.122 | 0.094 |
|  |  | 2084 | 0.111 | 0.092 |
|  |  | 3695 | 0.128 | 0.099 |
|  |  | 4761 | 0.120 | 0.095 |
|  |  | 3991 | 0.131 | 0.092 |

<sup>a</sup> SPT movies were recorded for 30 s total observation period at 20 Hz acquisition frame rate.

**Table S6.**
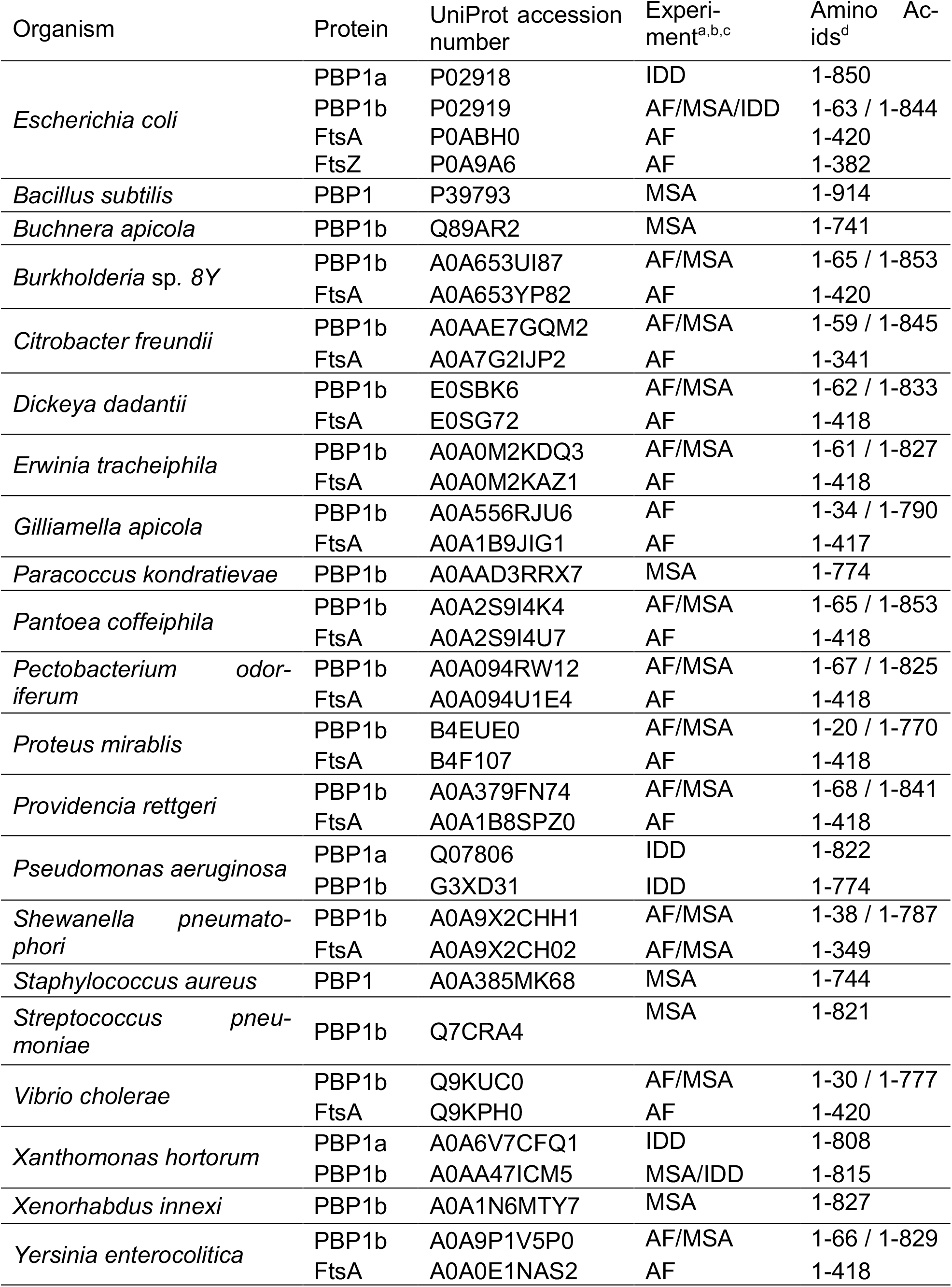
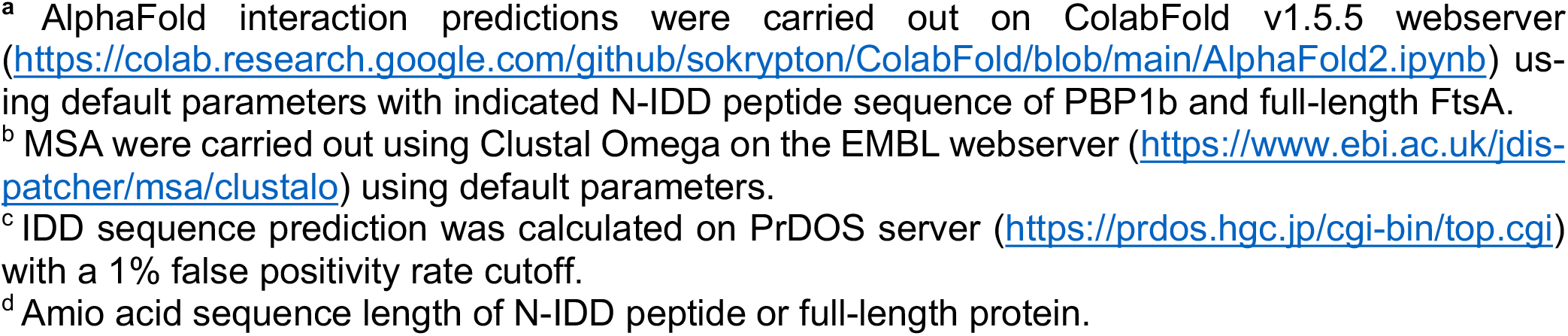
UniProt accession number of tested proteins for *in silico* analyses.

| Organism | Protein | UniProt accession number | Experiment<sup>a,b,c</sup> | Amino Acids<sup>d</sup> |
| --- | --- | --- | --- | --- |
| *Escherichia coli* | PBP1a | P02918 | IDD | 1-850 |
|  | PBP1b | P02919 | AF/MSA/IDD | 1-63 / 1-844 |
|  | FtsA | P0ABH0 | AF | 1-420 |
|  | FtsZ | P0A9A6 | AF | 1-382 |
| *Bacillus subtilis* | PBP1 | P39793 | MSA | 1-914 |
| *Buchnera apicola* | PBP1b | Q89AR2 | MSA | 1-741 |
| *Burkholderia* sp. 8Y | PBP1b | A0A653UI87 | AF/MSA | 1-65 / 1-853 |
|  | FtsA | A0A653YP82 | AF | 1-420 |
| *Citrobacter freundii* | PBP1b | A0AAE7GQM2 | AF/MSA | 1-59 / 1-845 |
|  | FtsA | A0A7G2IJP2 | AF | 1-341 |
| *Dickeya dadantii* | PBP1b | E0SBK6 | AF/MSA | 1-62 / 1-833 |
|  | FtsA | E0SG72 | AF | 1-418 |
| *Erwinia tracheiphila* | PBP1b | A0A0M2KDQ3 | AF/MSA | 1-61 / 1-827 |
|  | FtsA | A0A0M2KAZ1 | AF | 1-418 |
| *Gilliamella apicola* | PBP1b | A0A556RJU6 | AF | 1-34 / 1-790 |
|  | FtsA | A0A1B9JIG1 | AF | 1-417 |
| *Paracoccus kondratievae* | PBP1b | A0AAD3RRX7 | MSA | 1-774 |
| *Pantoea coffeiphila* | PBP1b | A0A2S9I4K4 | AF/MSA | 1-65 / 1-853 |
|  | FtsA | A0A2S9I4U7 | AF | 1-418 |
| *Pectobacterium odoriferum* | PBP1b | A0A094RW12 | AF/MSA | 1-67 / 1-825 |
|  | FtsA | A0A094U1E4 | AF | 1-418 |
| *Proteus mirabilis* | PBP1b | B4EUE0 | AF/MSA | 1-20 / 1-770 |
|  | FtsA | B4F107 | AF | 1-418 |
| *Providencia rettgeri* | PBP1b | A0A379FN74 | AF/MSA | 1-68 / 1-841 |
|  | FtsA | A0A1B8SPZ0 | AF | 1-418 |
| *Pseudomonas aeruginosa* | PBP1a | Q07806 | IDD | 1-822 |
|  | PBP1b | G3XD31 | IDD | 1-774 |
| *Shewanella pneumatophori* | PBP1b | A0A9X2CHH1 | AF/MSA | 1-38 / 1-787 |
|  | FtsA | A0A9X2CH02 | AF/MSA | 1-349 |
| *Staphylococcus aureus* | PBP1 | A0A385MK68 | MSA | 1-744 |
| *Streptococcus pneumoniae* | PBP1b | Q7CRA4 | MSA | 1-821 |
| *Vibrio cholerae* | PBP1b | Q9KUC0 | AF/MSA | 1-30 / 1-777 |
|  | FtsA | Q9KPH0 | AF | 1-420 |
| *Xanthomonas hortorum* | PBP1a | A0A6V7CFQ1 | IDD | 1-808 |
|  | PBP1b | A0AA47ICM5 | MSA/IDD | 1-815 |
| *Xenorhabdus innexi* | PBP1b | A0A1N6MTY7 | MSA | 1-827 |
| *Yersinia enterocolitica* | PBP1b | A0A9P1V5P0 | AF/MSA | 1-66 / 1-829 |
|  | FtsA | A0A0E1NAS2 | AF | 1-418 |

- <sup>a</sup> AlphaFold interaction predictions were carried out on ColabFold v1.5.5 webserver (https://colab.research.google.com/github/sokrypton/ColabFold/blob/main/AlphaFold2.ipynb) using default parameters with indicated N-IDD peptide sequence of PBP1b and full-length FtsA. - <sup>b</sup> MSA were carried out using Clustal Omega on the EMBL webserver (https://www.ebi.ac.uk/jdispatcher/msa/clustalo) using default parameters. - <sup>c</sup> IDD sequence prediction was calculated on PrDOS server (https://prdos.hgc.jp/cgi-bin/top.cgi) with a 1% false positivity rate cutoff. - <sup>d</sup> Amino acid sequence length of N-IDD peptide or full-length protein.

**Video S1: *In situ* architecture of sPG in wild-type *E. coli***. Cryo-electron tomogram of wild-type *E. coli*. Time-lapse series were acquired with a rate of 7 fps in the compressed format m4v for visualization purposes. Green, cyan and magenta layers indicate segmented IM, PG and OM, respectively. Ribosomes are shown in yellow. Scale bars = 100 nm.

**Video S2: *In situ architecture of sPG in* Δ*ponB E. coli***. Cryo-electron tomograms of Δ*ponB* cells. Details as for Video S1.

**Video S3: *In situ architecture of sPG in* Δ*lpoB E. coli***. Cryo-electron tomograms of Δ*lpoB* cells. Details as for Video S1.

**Video S4: *In situ architecture of sPG in* Δ*ponA E. coli***. Cryo-electron tomograms of Δ*ponA* cells. Details as for Video S1.

**Video S5: Time-lapse video of *E. coli* cells subjected to osmotic oscillations**. Indicated strains were imaged in a microfluidic flow cell (CellAsic) in presence of 1µM SytoxGreen or 1µM propidium iodide (^α^*ponB(R6E)*). Osmotic oscillations were performed by switching media from LB to 0.5xLB0N ten times over a 42min observation period. Images were acquired at 0.1 Hz. Individual phase (center) and fluorescence (right) channels are shown in addition to a merged overlay (left). Representative examples for WT, Δ*ponB*, Δ*ponB pNP146 (P*_*ara*_::*sulA)*, Δ*ponB pAV39 (P*_*lac*_::*ponB(R6E, M46L))*, Δ*lpoB, and* Δ*ponA* are sequentially shown. For further details, see *Methods*. Scale bar = 2 µm.

**Video S6: Side-by-side comparison of SPT experiments with *E. coli* cells expressing Halo fusions to the indicated PBP1b isoforms**. STP trajectories of indicated Halo-PBP1b isoforms (labeled with 20 nM JF549) are colored according to their mean track speed (blue = slow, red = fast). ZapA-sfGFP (false-colored in green) was used as a fiducial marker for the divisome and overlayed over a bright field reference image. Images were recorded at 20 Hz and binned 2×2. For further details, see *Methods*. Scale bar = 0.5 µm.

**Video S7: FtsZ treadmilling assay in response to N-**^**α**^**PBP1b peptide addition**. Alexa488 labeled FtsZ (1.25 µM) and unlabeled FtsA (0.4 μM) were reconstituted on supported lipid bilayers (SLBs) in the presence of 4 mM ATP/GTP and allowed to self-organize into treadmilling filaments and followed by TIRF microscopy at 0.5 Hz acquisition frame rate. At 2 min, 0.4 µM Cy5 labled N-^α^PBP1b peptide was added to the reaction chamber. For further details, see *Methods*. Scale bar = 5 µm.

**Video S8: FtsZ treadmilling assay in response to N-**^**α**^**PBP1b(R6E) peptide addition**. Alexa488 labeled FtsZ (1.25 µM) and unlabeled FtsA (0.4 μM) were reconstituted on SLBs in presence of 4 mM ATP/GTP and allowed to self-organize into treadmilling filaments and followed by TIRF microscopy at 0.5 Hz acquisition frame rate. At 2 min, 0.4 µM Cy5 labled N-^α^PBP1b(R6E) peptide was added to the reaction chamber. For further details, see *Methods*. Scale bar = 5 µm.

**Extended Data Table 1**. Hits from AlphaFold multimer screen for PBP1b isoforms. For additional information check: https://private.predictomes.org/library/help.

